# Clonal selection of somatic mutations shapes NK cell memory

**DOI:** 10.64898/2026.09.17.752421

**Authors:** Timo Rückert, Matthias Günther, Oliver Knight, Maximilian Mandry, Karl Junker, Simon Grassmann, Joseph C. Sun, Thomas Höfer, Chiara Romagnani

## Abstract

Adaptive immune memory relies on selective clonal expansion and persistence of lymphocyte populations. Whether and how clonal selection occurs in innate lymphocytes that lack rearranged antigen receptors remains poorly understood. Here, we used somatic genetics to reconstruct the natural history of human natural killer (NK) cell responses to cytomegalovirus infection. Across a large cohort of healthy seropositive individuals, we found strong clonal dominance in memory NK cells consistent with clonal selection. Dominant memory NK cell clones were marked by coherent receptor and chromatin configurations, emerged early in life and diversified into genetically related subclones. Both an excess of coding mutations, and the reconstructed population dynamics of the dominant clones, provided evidence for positive selection acting on somatic genetic variation. We identified somatic loss-of-function mutations in the chromatin modifier *KMT2D*, and showed in a mouse model that *Kmt2d* deletion alters NK-cell fitness depending on timing and environmental context. Together, these findings establish clonal selection as an organizing principle of human NK-cell memory and support a model in which heritable cell states and acquired genetic variation contribute to shape clonal competition.

---

Clonal selection is a defining principle of adaptive immunity: rare antigen-specific lymphocytes expand into large effector populations and can establish long-lived immunological memory. For T and B cells, antigen receptor affinity and naive precursor frequency are among the main determinants of clonal burst size and differentiation^1,2^, despite the fundamentally stochastic nature of individual antigen-specific T cell responses^3,4^. Whether analogous selection operates among innate lymphocytes, and which mechanisms might underlie clonal selection in the absence of somatically diversified antigen receptors, remains unclear.

Somatic mutations provide a unique means of investigating these processes by serving as endogenous lineage barcodes, allowing clonal relationships and cellular histories to be reconstructed directly in vivo^5–7^. While most acquired mutations are selectively neutral, some alter cellular fitness and become enriched by driving positive selection. In the hematopoietic system, population-scale sequencing studies have identified recurrent driver mutations underlying clonal hematopoiesis, an age-associated expansion of blood-cell clones frequently involving genes implicated in myeloid malignancies. More recently, genetic signatures of positive selection have also been detected in T and B lymphocytes from healthy individuals, demonstrating that somatic variation outside antigen-receptor loci can influence fitness even of naive lymphocytes^8,9^.

Persistent or latent viral infection provides a physiological context in which these forces can be investigated in lymphocytes. Cytomegalovirus (CMV) drives extensive expansions of virus-specific CD4^+^ and CD8^+^ T cells, progressively reshaping T cell receptor repertoires and producing pronounced oligoclonal dominance, indicative of strong selection pressures^10–12^. CMV also induces durable adaptations in natural killer (NK) cell subsets through a distinctive combination of inflammatory signals and receptor–ligand interactions, including direct recognition of the viral m157 protein by the Ly49H receptor in mice^13^ and CD94/NKG2C-HLA-E-dependent responses to peptides from the viral UL40 protein in humans^14–17^. Both mouse^18^ Ly49H^+^ and human NKG2C^+^ NK cells can undergo clonal expansion in response to CMV. Expanded NK cell clones persist with clone-associated epigenetic states in healthy human CMV (HCMV)-seropositive individuals^19^, giving rise to long-lasting memory NK cells, commonly referred to as adaptive NK cells^20^ or NK3^21^. These findings have established the clonal maintenance of NK-cell memory but left unresolved the extent of clonal dominance, the underlying cell population dynamics and, crucially, whether the clonal architecture is shaped by selection.

Here, we combine somatic mutation profiling, single-cell phylogenies and mutation-based modelling to reconstruct the evolutionary history of HCMV-associated memory NK-cell expansions across a large cohort of healthy individuals. We uncover an unexpectedly large degree of clonal dominance of memory NK cells that develops at an age consistent with HCMV infection. Orthogonal population-genetic analyses strongly indicate that clonal dominance arises by selection. Supporting this conclusion, we identify two predicted loss-of-function variants in the epigenetic regulator *KMT2D* and demonstrate in a mouse model that Kmt2d loss alters NK-cell competitive fitness, with the direction of selection determined by cellular state and the timing of deletion.

## Massive NK-cell clones dominate the memory compartment in HCMV-seropositive individuals

To quantify the clonality of HCMV-associated NK-cell memory at scale, we recruited a cohort of 81 healthy HCMV-seropositive individuals. Adaptive NK cell expansions were identified by flow cytometry (Methods) and covered a wide range of adaptive NK cell population sizes (Extended Data Fig. 1a-b). We performed whole-exome sequencing (WES) of sorted NKG2C^+^ memory NK cells (N=81) and paired conventional NK cells (N= 15), with monocytes serving as a control for removal of germline variants in both populations (Methods and Fig. 1a). In contrast to conventional NK cells, memory NK cells frequently harbored numerous high variant allele frequency (VAF) mutations (Fig. 1b–c, Extended Data Fig. 1c). The high-VAF mutations detected in memory NK cells jointly formed discrete VAF peaks in many individuals, suggesting that co-occurring mutations were carried by large memory clones. Corresponding peaks in the paired conventional compartment were absent, consistent with its polyclonality (Fig. 1c, Extended Data Fig. 1c).

**Figure 1.**
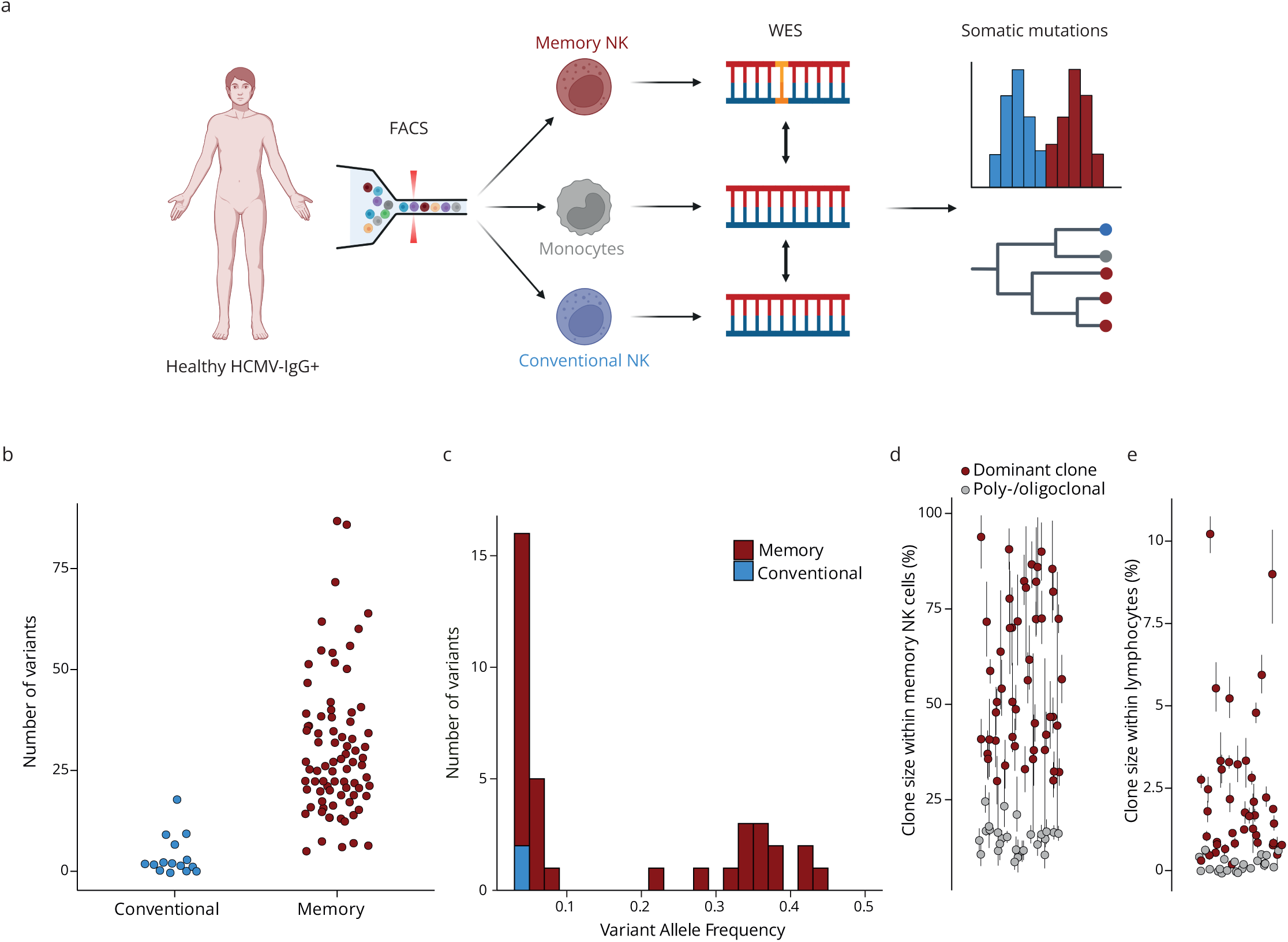
Dominant memory NK-cell clones are widespread in healthy HCMV-seropositive individuals. **a** Sorted memory (NKG2C^+^ NKG2A^-^, N = 81, 114x median coverage) and conventional (NKG2C^-^ NKG2A^+^ CD57^-^, N= 15, 126x median coverage) NK cells were analyzed by WES, using monocytes (CD14^+^, 61x median coverage) as a germline reference to call somatic mutations for estimation of clone size distributions and phylogenies. Created with Biorender.com. **b** Number of somatic mutations at VAF ≥5% in conventional and memory NK cells. **c** Representative VAF histogram of NK cell subsets from donor HC02. **d-e** Leading memory NK cell clone size estimates with 95% confidence intervals from SCIFER^7^ within the memory NK cell (**d**, N = 76) and lymphocyte compartments (**e**, N = 66).

We estimated the fraction of memory NK cells belonging to the leading clone using SCIFER^7^ for all individuals with at least 10 identified variants (N = 76) (Methods, Extended Data Fig. 2a). Using a conservative threshold of >25%, we observed that 49 of 76 individuals (64%) carried a dominant clone occupying more than 25% of the NKG2C^+^ memory compartment, with the largest reaching 95% (median 39%) (Fig. 1d, Extended Data Fig. 2b). These clone size estimates captured a much larger fraction of the memory NK cell compartment compared to previously published^19^ and newly generated data analyzing mitochondrial clonotypes, which underestimated clonal dominance even when considering their total cumulative fraction (Extended Data Fig. 2c), underlining the superiority of the present approach for faithful quantification of NK cell clonality.

To quantify the size of these clones beyond the memory NK-cell compartment, we integrated inferred clone frequencies with immunophenotyping by flow cytometry (N = 66). A substantial fraction of the total lymphocyte pool was derived from dominant memory NK cell clones, accounting for up to 10% (median 0.9%) of all circulating lymphocytes (Fig. 1e, Extended Data Fig. 2d). The frequency of the leading clone was not detectably correlated with the proportion of NKG2C^+^ cells (Extended Data Fig. 2e), indicating that greater clonal dominance was not consistently accompanied by enlargement of the memory NK cell compartment.

Together, these data establish that the HCMV-associated NK-cell memory compartment is characterized by widespread and unexpectedly pronounced clonal dominance, which can substantially reshape the clonal composition and somatic mosaicism of the circulating lymphocyte pool.

## Extensive branching in clonal phylogenies links expansion to stable NK-cell states

Having quantified the prevalence and magnitude of clonal dominance within the memory NK cell compartment, we next sought to jointly resolve NK cell clonal architectures and corresponding surface phenotypes in an unbiased manner. To this end, we designed donor-specific panels to perform joint single-cell profiling of somatic variants and surface proteins in total NK cells for three representative individuals (Methods). This analysis identified 2-5 independent clonal families per donor defined by co-occurring somatic mutations (Fig. 2a,b, Extended Data Fig. 3a,b). We found a range of differently sized clonal families that individually comprised 2.3-63% of the total NK cell pool (Extended Data Fig. 3c) or up to 7% of lymphocytes, which is equivalent to ∼7 x 10^8^ circulating cells assuming average lymphocyte counts^22^ (Fig. 2c). Leading-clone frequencies within the NKG2C^+^ memory compartment were highly concordant between single-cell and bulk sequencing (Extended Data Fig. 3d), independently validating our clone size estimates. Reconstruction of subclonal phylogenies revealed prominent branching architectures consistent with substantial proliferative history during which subclonal mutations were acquired (Fig. 2a,b, Extended Data Fig. 3a,b). These individual clonal families displayed up to five internal nodes (Fig. 2b) supported by 11-16 co-occurring variants in the most expanded clones (Fig. 2a, Extended Data Fig. 3a,b). Thus, the dominant clones detected by bulk sequencing encompassed genetically diversified descendants of a single founder cell.

**Figure 2.**
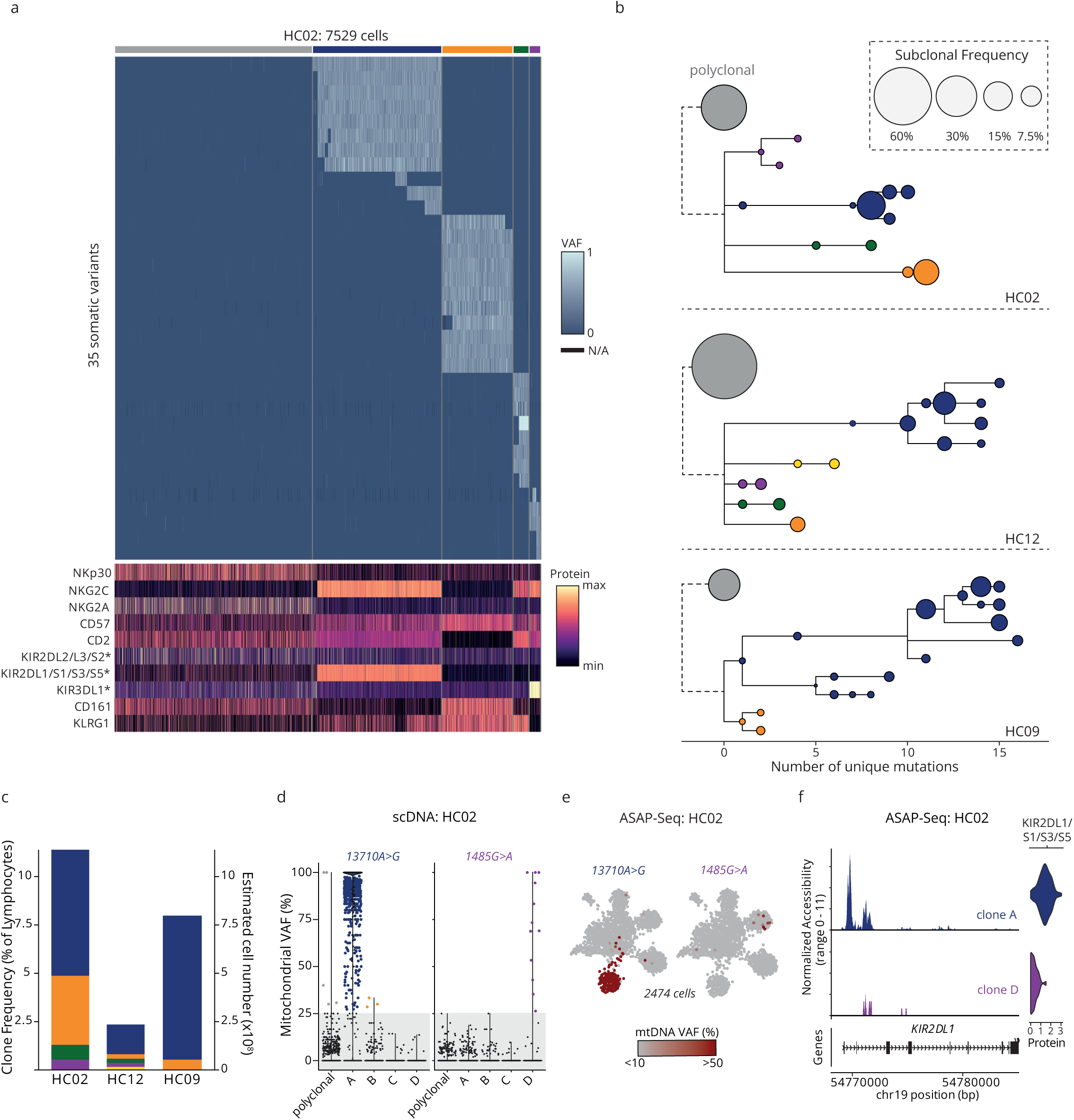
Extensive branching in clonal phylogenies links expansion to stable NK-cell states. **a** Joint single-cell profiling of cell surface antigens and somatic mutations in donor HC02 (scDNA-seq). Asterisks mark self-MHC-specific KIRs. **b** Reconstructed clonal phylogenies with subclonal frequencies derived from single-cell mutational profiling. **c** Proportions of individual clones within total lymphocytes and estimation of absolute cell numbers. **d-e** Clone-associated mtDNA variants in scDNA- (**d**) and ASAP-seq (**e**) for donor HC02. **f** Clone-specific chromatin accessibility profiles at the clonally segregated *KIR2DL1* locus, shown alongside surface protein levels quantified by ASAPseq.

Simultaneous unbiased profiling of cell surface proteins validated that most expanded clonal families identified in these individuals exhibited a characteristic memory NK-cell phenotype, NKp30^lo^NKG2C^+^NKG2A⁻CD2^+^, although clones with non-canonical phenotypes were also observed (Fig. 2a, Extended Data Fig. 3a,b). Cells lacking detected clone-defining mutations were phenotypically diverse, as expected for conventional NK cells, whereas those within individual clonal families displayed highly restricted NK receptor-expression profiles (Fig. 2a, Extended Data Fig. 3a,b). Even within the same individual, distinct clonal families frequently expressed different, and within each family coherent, combinations of self-MHC-specific killer-cell immunoglobulin-like receptors (sKIRs) (Fig. 2a, Extended Data Fig. 3a,b). These differences occurring even within a shared host environment were consistent with clonal propagation of single mature NK cells characterized by distinct receptor states.

Building on our previous observation of clone-associated epigenetic states^19^, as assessed by combining analysis of mitochondrial mutations with ASAP-seq^19,23^, we next asked how receptor configurations related to chromatin accessibility within genetically defined clonal families. To this end, we included mitochondrial DNA (mtDNA) mutations in our targeted single cell genotyping panel and used those detectable in both assays to map mitochondrial clonotypes onto clonal families identified by nuclear variants and vice versa (Fig. 2d,e, Extended Data Fig. 3e,f). Apart from one large mtDNA clonotype in HC02 (*13710A>G*) that marked virtually all cells of clone A (Fig. 2d,e), the other detected clonotypes were subclonal (Fig. 2d,e, Extended Data Fig. 3e,f), potentially explaining the observed discrepancy in clone size estimates (Extended Data Fig. 2c). Clonotypes linked to different clonal families occupied distinct chromatin accessibility states (Fig. 2e, Extended Data Fig. 3f). In particular, clone-specific accessibility at KIR promoters corresponded closely to expression of the respective surface receptors, linking clonal receptor configurations to their underlying regulatory states (Fig. 2f, Extended Data Fig. 3g).

Together, these findings resolve the genetic heterogeneity of massively expanded memory NK cell clones. Further acquisition of subclonal variants during expansion results in branching clonal architectures that raise questions on clonal timing and expansion dynamics. Conversely, expanded clonal families retain coherent receptor and regulatory states despite internal genetic diversification. Such clonally propagated differences provide candidate phenotypic substrates for selection, while the unequal abundance of clonal families shapes the population-level NK-cell receptor repertoire.

## Memory NK cell clones emerge at a young age and undergo slow population growth

The striking dominance and size of memory NK cell clones prompted us to next ask when the founding cells first emerge, and how fast the clones grow. To this end, we used a population-genetics approach that quantifies the statistics of neutral somatic single nucleotide variants (SNVs) emerging during clonal growth from whole-genome sequencing (WGS) data (Methods)^7,24^. WGS data of memory NK cells from the same three individuals analyzed by single cell mutation profiling (aged 28, 29 and 49 years) demonstrated that mutations were mainly acquired from clock-like mutational processes^25^ manifesting in single base substitution (SBS) signatures SBS1, and the highly similar SBS5 and SBS40a (Fig. 3a,b).

**Figure 3.**
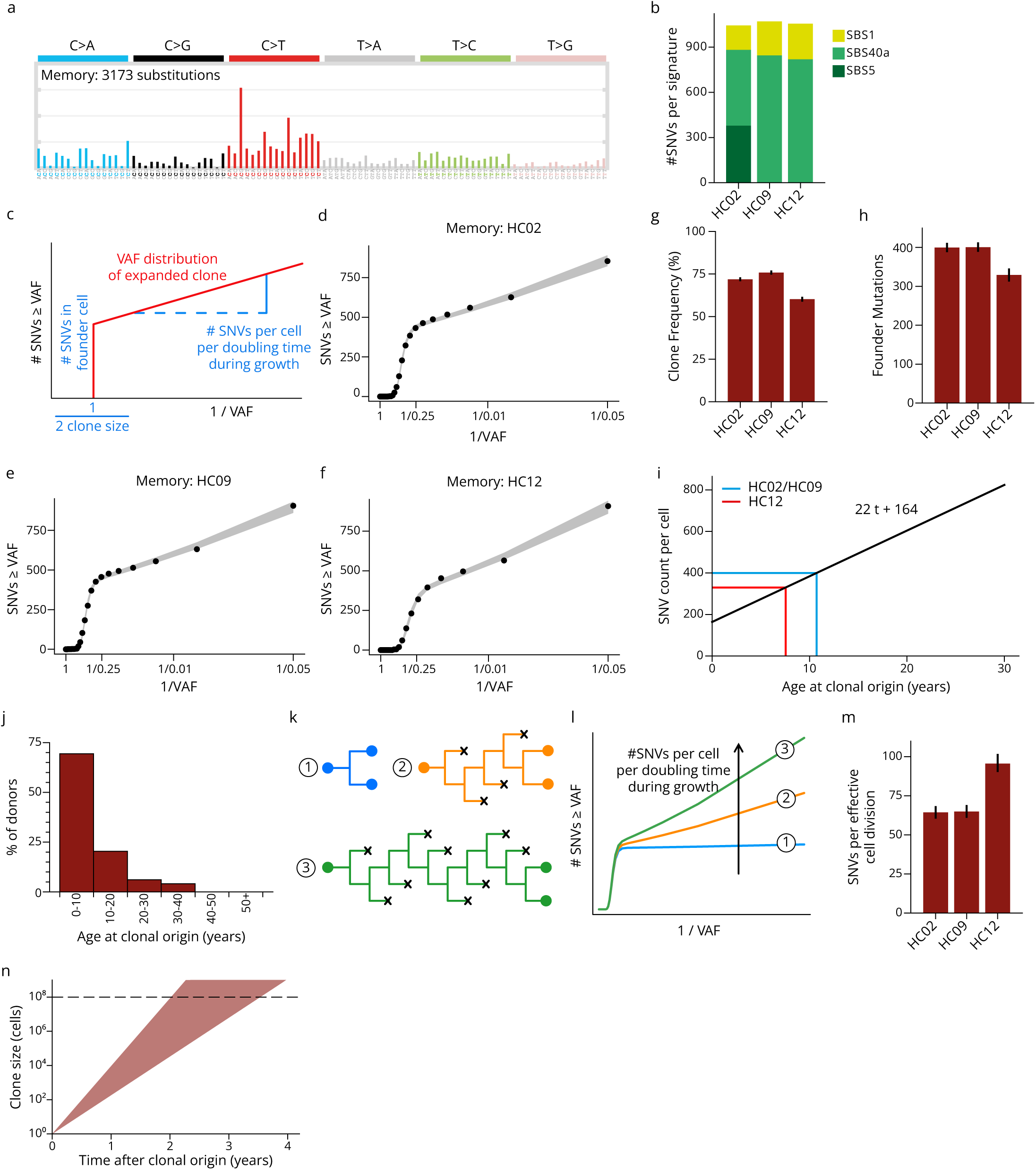
Memory NK cell clones emerge at a young age and undergo slow population growth. **a** Genome-wide somatic substitution pattern aggregated over three donors and **b** quantified as SBS signatures. **c** Schematic of population-genetic analysis of the VAF histogram for each individual donor. **d-f** VAF histograms of three donors. **g** Clone size and **h** Number of SNVs in the founding cell of each leading clone inferred from the VAF histograms. **i** Real-timing of the donor’s age at which the clonal founding cell arose for the three donors subjected to WGS. **j** Real-timing of the age of the emergence of the clonal founding cell across the individuals in the WES cohort (N = 49). **k** Schematic showing that a variable number of cell divisions (blue, 1; orange, 5; green, 10) might occur for one doubling event (x, cell loss by differentiation into a non-dividing state or cell death). **l** Different modes of cell doubling (shown in **k**) shape the subclonal tail of the VAF histogram. **m** Number of SNVs acquired per cell doubling as inferred from subclonal tails of the VAF histograms. **n** Predicted clonal growth time at different sizes and for inferred effective division rates (shaded).

Clonality causes a shoulder in the cumulative histogram of VAFs^7^ (Fig. 3c), which is evident for all three memory NK cell samples (Fig. 3d-f). Moreover, the VAF histogram yields three key parameters (Fig. 3c): (i) the clone size, (ii) the number of SNVs that accumulated in the clonal founding cell, which reflects the age at which the founding cell emerged, and (iii) the number of SNVs that were acquired per doubling time of the clone size during clonal growth. The clone sizes determined by WGS matched those estimated by both WES and targeted sequencing (Fig. 3g, Extended Data Fig. 3d). Remarkably, the number of SNVs in the clonal founding cells was similar in all individuals, despite individual HC02 being ∼20 years older than the other two (Fig. 3h), suggesting that the clones emerged at similar ages in all three donors. To real-time clone origins from the number of founder mutations, we used accurate measurements of SNV accumulation in naive T lymphocytes across the human lifespan^9^. For the individuals at study, this yields clone origins at young age, between 7 and 11 years of age (Fig. 3i). We then extrapolated these findings to the individuals with dominant clones in the WES cohort (Methods). Nearly 70% of single dominant clones were estimated to have emerged within the first ten years of life, approximately 20% between 10 and 20, and the remainder at an older age (Fig. 3j), broadly coinciding with the age range of HCMV-seroconversion in the German population^26^.

Finally, we asked how rapidly the memory NK cell clones grew to their size of ∼10^8^ cells (Fig. 2c). Based on Ki67 staining of NKG2C^+^ NK cells during infection, the proliferation rate in the acute expansion phase can be large, with 30-70% of cells being in cycle at any one time^27^. Theoretically, if all NK cells contributed to population expansion (Fig. 3k, blue cell division, leading to two further dividing daughter cells) with an acute proliferation rate of once per day, a clone could grow to size 10^8^ in about two and a half weeks. However, clonal expansion could be carried by only a subset of cells, if differentiation into a non-dividing state or/and cell death also occurred (Fig. 3k, orange and green division cell pedigrees, with different degrees of loss of proliferating cells by differentiation or death). The subclonal tail of the cumulative VAF histogram measures the degree of cell loss during clonal expansion (Fig. 3l)^7,24^. Indeed, we find that 60-100 SNVs accumulate for each cell doubling in the expanding memory NK populations of all three donors (Fig. 3m). Given that about 1.5 SNVs typically occur per cell division^7^, this suggests that only one in 40-70 NK cell divisions are self-renewing and hence contribute to clonal growth, which is consistent with the extensive subclonal branching we observed in single cell genotyping (Fig. 2b). As a consequence, we predict growth times of the memory NK cell clones of several years after the founding event (Fig. 3n).

## Somatic mutations in memory NK cells undergo clonal selection

The magnitude and prevalence of dominant clones within the memory NK compartment prompted us to ask whether these expansions could plausibly arise through neutral drift. The efficiency of drift strongly depends on population size and the number of cellular generations. NK cells expressing the HCMV-associated receptor combination NKG2C^+^ NKG2A^-^ CD2^+^ sKIR^+^ already comprise approximately 1% of the NK cell compartment in HCMV-seronegative individuals^14,28,29^, corresponding to ∼10⁷ circulating cells^22^. Moreover, clonal expansion was not restricted to this receptor configuration (Fig. 2a, Extended Data Fig. 3a,b), suggesting a larger pool of cells can participate in the response. However, even assuming a substantially smaller responsive population of only ∼10⁵ cells and an exceptionally rapid steady-state division rate of once per day, neutral drift would require more than a century to generate comparable clonal dominance (Fig. 4a, Methods). At more realistic, substantially lower turnover rates, the expected timescale increases further. Thus, neutral drift within a large, stably maintained NK-cell population was insufficient to explain the observed clonal architecture.

**Figure 4.**
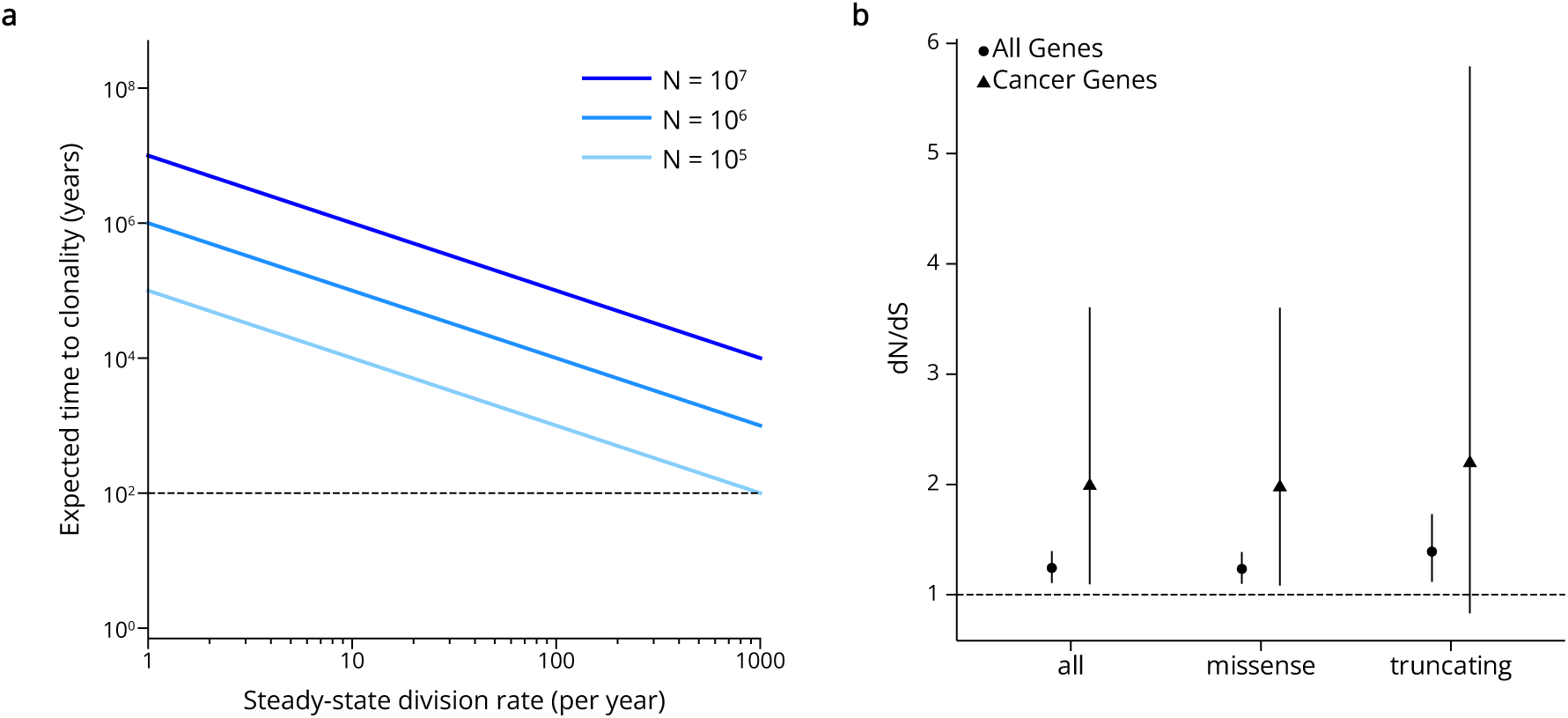
Evidence of clonal selection from population dynamics and genetic signatures. **a** Neutral drift time estimates at different population sizes and steady-state division rates. **b** dN/dS scores with 95% confidence intervals for all genes and cancer-associated genes (Cancer Gene Census Tier 1).

We next asked whether acquired coding mutations are under selective pressure. We therefore analyzed the ratio of nonsynonymous to synonymous mutations using a model that accounts for sequence-context-dependent mutation rates^30^. Across the exome, nonsynonymous substitutions were enriched relative to the neutral expectation (dN/dS = 1.24, 95% confidence interval, 1.11-1.39; Fig. 4b), providing evidence of positive selection on acquired coding variation, with 1 in 5 (range 1 in 10 to 1 in 4) non-synonymous mutations conferring a selective advantage.

Although the cohort of 81 donors analyzed afforded limited power to identify individual genes under selection, analysis of cancer-associated genes^31^ also detected a positive-selection signal (dN/dS = 1.98, 95% confidence interval, 1.09–3.61, Fig. 4b). Among those, we identified several potentially functional mutations in genes previously implicated in lymphocyte or NK-cell biology, including truncating variants as well as missense substitutions with high AlphaMissense^32^ (AM) scores. These included a nonsense variant in *LEF1*, which is recurrently inactivated in T-cell acute lymphoblastic leukemia^33,34^, and a frameshift variant in *TNFAIP3* (p.Glu88AlafsTer7), encoding the negative NF-κB regulator A20, which is frequently inactivated in lymphomas^35,36^. We further identified a missense variant in the DNA-binding domain of *PRDM1* (p.Thr626Lys; AM score = 0.9335), a transcriptional repressor recurrently inactivated in NK-cell malignancies^37^. Notably, we also detected a previously reported gain-of-function variant in *STAT3* (p.Asn647Ile), located within the SH2-domain mutational hotspot recurrently affected in NK- and CD8^+^ T-cell lineage large granular lymphocytic leukemia^38^. These alterations nominated candidate mechanisms through which somatic variation could influence NK-cell fitness, although the aggregate selection signal did not establish the contribution of each individual variant.

Together, population modelling and coding-mutation spectra support selection as a contributor to memory NK-cell clonal dominance, with acquired genetic variation as one substrate. These findings motivated functional testing of a candidate regulator to determine how its disruption affects NK-cell competitive fitness.

## Kmt2d loss affects NK-cell competitive fitness

We next asked whether disruption of a gene identified through human somatic mutation profiling could alter NK-cell competitive fitness. We focused on the chromatin modifier *KMT2D*, in which we identified two independent mutations in memory NK cells from distinct individuals: a nonsense variant, p.Arg2687Ter, predicted to cause loss of function through nonsense-mediated decay^39^, and a missense variant, p.Tyr1495Asp, with a high predicted functional impact (AM score 0.9963). *KMT2D* is recurrently affected by loss-of-function mutations in B-cell malignancies^40,41^, and mutations have been reported in clonal hematopoiesis^8^ as well as NK-cell lymphoproliferative disorders^42^, while its role in NK cell biology has not been explored. These observations prompted us to investigate how constitutive or inducible Kmt2d deficiency influences NK-cell competition in a mouse model at steady-state or during infection.

To this end, we generated mice with conditional *Kmt2d* deletion in NK cells (*Ncr1-Cre Kmt2d-flox*) and all hematopoietic cells (*Vav1-Cre Kmt2d-flox*). At steady state, Kmt2d deficiency increased NK-cell frequencies in a gene-dose-dependent manner (Extended Data Fig. 4a–c). Consistently, Kmt2d-deficient NK cells progressively increased in relative abundance compared with wild-type competitors in mixed bone marrow chimeras, demonstrating a competitive advantage (Fig. 5a–c).

**Figure 5.**
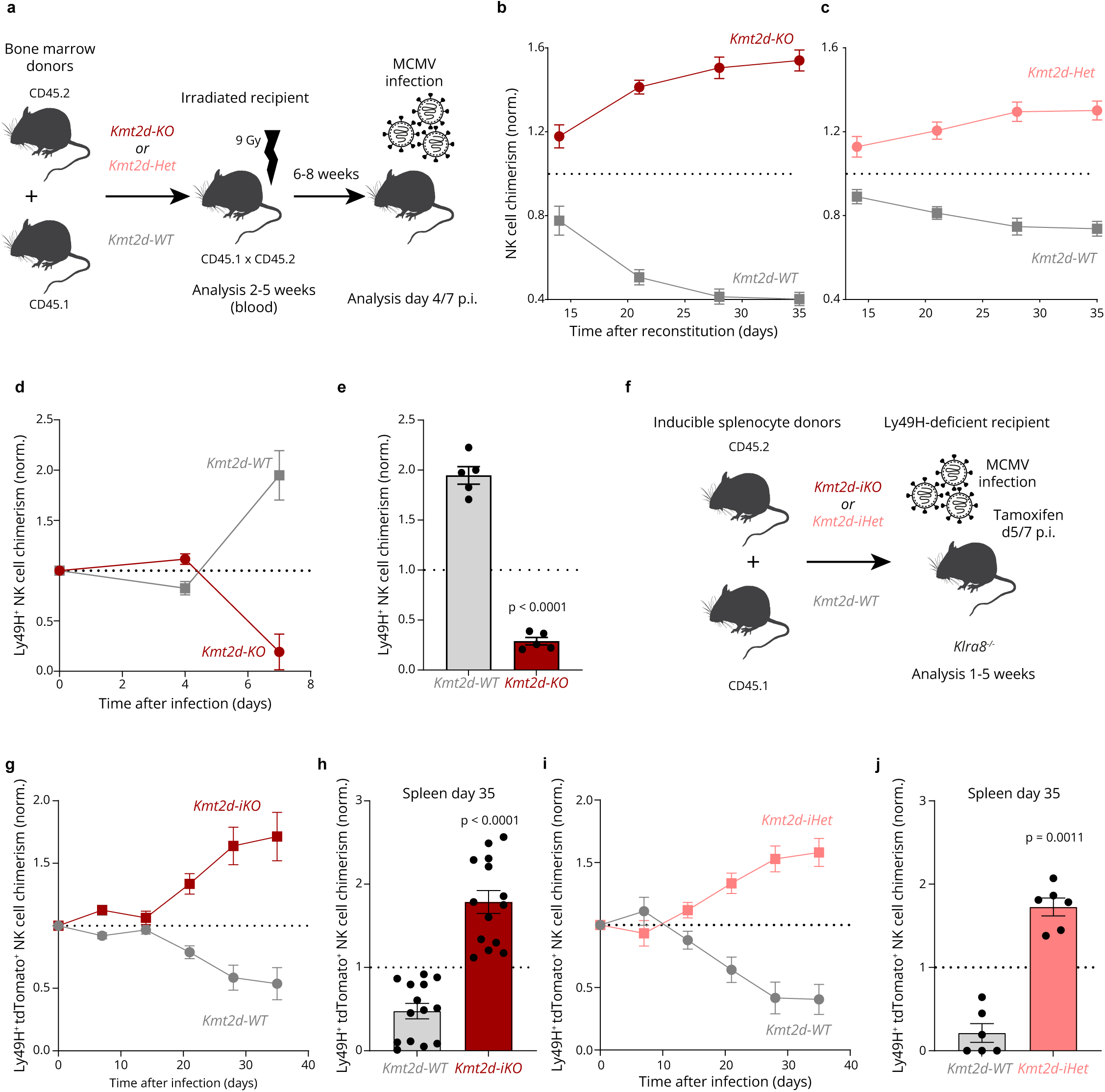
*Kmt2d* loss-of-function is subject to context-dependent selection in vivo. **a** Congenically marked hosts were irradiated and reconstituted with NK-cell-depleted bone marrow cells from *Ncr1^Cre^*x *Kmt2d^fl/fl^*(Kmt2d-KO) or *Ncr1^Cre^*x *Kmt2d^fl/wt^*(Kmt2d-Het) mixed with *Ncr1^Cre^*x *Kmt2d^wt/wt^*(WT) controls. After 6-8 weeks, mixed bone marrow chimeras were infected with MCMV. **b-c** Reconstitution of Kmt2d-KO (N=9-10) (b) and Kmt2d-Het (N=5) (c) NK cells in the blood of mixed bone marrow chimeras, frequencies normalized to lineage-positive cells. **d-e** Relative expansion in blood (d) and spleen (e) of Kmt2d-ko NK cells after infection of mixed bone marrow chimeras with MCMV (N=5-8). **f** *Ubc^CreERT2^* x *Kmt2d^fl/fl^* x *tdTomato^LSL^* (Kmt2d-iKO) or *Ubc^CreERT2^* x *Kmt2d^fl/wt^* x *tdTomato^LSL^* (Kmt2d-iHet) splenocytes were co-transferred with *Ubc^CreERT2^*x *Kmt2d^wt/wt^*x *tdTomato^LSL^*(WT) into Klra8^-/-^ (encoding Ly49H) hosts before infection with MCMV. At the peak of infection (day 5-7) mice were treated with tamoxifen to induce *Kmt2d*-deletion. **g-j** Relative chimerism of Kmt2d-iKO (N=12-15) (g,h) and Kmt2d-iHet (N=5-10) (i,j) in blood (g,i) and spleen (h,j) after MCMV infection. Summary graphs display mean with standard error, statistical significance from one-sample t-test against normalized chimerism of 1.

We then assessed how Kmt2d-deficiency affected NK cell responses to murine CMV (MCMV) infection. In contrast to the competitive advantage observed at steady-state, Kmt2d-deficient Ly49H⁺ NK cells expanded less effectively than Kmt2d-sufficient competitors (Fig. 5d,e), and exhibited altered expression of maturation markers CD27 and CD11b (Extended Data Fig. 4d,e). In a competitive adoptive-transfer setting, impaired acute expansion was followed by a sustained disadvantage at later time points (Extended Data Fig. 4f–h).

As these observations suggested a negative effect when Kmt2d is lost before infection and our retrospective analyses of established clones from humans ex vivo cannot precisely establish when individual mutations arose relative to infection, we next set out to model somatic *KMT2D* loss in mature NK cells after infection. Using a tamoxifen inducible model (*Ubc^CreERT2^* x *Kmt2d^fl/fl^* x *tdTomato^LSL^*) in a competitive splenocyte-transfer setting, we induced deletion on days 5-7 after MCMV infection (Fig. 5f), coinciding with maximal Kmt2d expression during the response (Extended Data Fig. 4i). In this setting, tdTomato^+^ NK cells carrying inducible Kmt2d homozygous deletion gained a competitive advantage that became apparent from day 21 (Fig. 5g–h). Similar effects were observed also after heterozygous deletion (Fig. 5i–j). By day 35, these cells dominated the donor-derived splenic memory NK-cell population (Fig. 5h,j). These data show that *Kmt2d* loss-of-function occurring after infection provides a competitive advantage, sustaining NK cell maintenance.

Together, these experiments identify KMT2D as a regulator of NK-cell competitive fitness and demonstrate that the loss in mature NK cells after MCMV infection can confer a positive selective effect in vivo. The opposing outcomes connect a gene identified through human somatic variation to a broader principle of clonal selection: the competitive consequences of a genetic alteration depend on the timing and the conditions under which it acts.

Overall, these findings establish clonal selection as a driving force underlying lymphocyte population dynamics which acts on heritable traits beyond rearranged antigen receptors.

## Discussion

We have previously established the clonal maintenance of NK cell memory in response to HCMV^19^. In this study, we elucidate the magnitude, prevalence, dynamics, and drivers of NK cell clonal selection and provide a quantitative framework that reconstructs the evolutionary processes underlying clonal dominance. Together, these observations support clonal selection as an organizing principle of NK-cell memory and introduce somatic evolution as a physiological immune adaptation of the innate lymphocyte compartment.

Despite the lack of rearranged receptors, the magnitude and prevalence of NK cell clonal dominance observed in HCMV-seropositive individuals is remarkable. The extraordinary proliferative capacity of lymphocytes provides the basis for extensive expansion^44^ but does not itself explain why particular clones become dominant. The determinants of clonal outgrowth are likely to be multidimensional^45^, reflecting the combined contributions of surface receptor expression, cell state and acquired genetic variation. Recurrent combinations of receptors on dominant clones, such as high NKG2C, CD2, and sKIR, along with lack of NKG2A expression, likely reflect clonal selection for optimally balanced integration of activating and inhibitory signals at different stages of the response, starting from the initial recruitment. High expression of NKG2C^17,27^ recognizing HCMV UL40 peptides presented by HLA-E^16^ is reminiscent of avidity selection in T cells, and a similar mechanism has been shown to support the expansion of mouse Ly49H^+^ NK cells after MCMV infection^46^. Conversely, lack of NKG2A can likely be attributed to a negative selective effect due to its higher affinity to the same HLA-E peptide ligands, as compared to NKG2C^47^. Other recurrent inhibitory receptors such as sKIRs^14^ could either play a role in education^48^ before NK cells get activated or provide a survival advantage later on^49^. Beside surface receptors, broader clone-associated regulatory programs and epigenetically inherited traits could potentially modulate clonal fitness. Similar interclonally diversified but intraclonally coherent and heritable cell states have been observed for human T cells^50^, suggesting that regulatory-state diversity may provide a general layer of functional variation across different lymphocyte populations.

We show that somatic mutations add a genetically heritable dimension to this diversity. The net positive selection observed in memory NK cells suggests that approximately 20% of detected non-synonymous somatic variants confer a clonal advantage, which is in line with recent estimates from T and B lymphocytes^9^. Although the full driver landscape remains elusive, signatures of selection in cancer-associated genes and exemplary mutations in genes implicated in lymphoid malignancies such as *KMT2D*, *TNFAIP3*, *PRDM1, LEF1,* and *STAT3* point towards shared pathways to enhanced fitness. However, in contrast to malignancies or clonal hematopoiesis where mutations are the primary drivers and result in strong enrichments of coding mutations in relatively defined sets of genes^30,51^, we did not observe significant recurrence of genes affected by damaging mutations. Despite the comparably small size of our cohort, this argues against a small set of mutations with strong growth effects and instead suggests diverse pressures at different phases of the anti-viral response and mutations that enhance fitness through multiple mechanisms.

Our mouse experiments provide in vivo proof of principle that disruption of a gene identified through our human somatic mutation profiling can alter NK-cell competitive fitness, using the epigenetic regulator Kmt2d as an example. Although these models cannot unequivocally reconstruct selection of the *KMT2D*-mutated clones in human individuals, they provide important insights on the functional role of this gene, as well as on the context dependent effects of Kmt2d loss-of-function. Our observation of reduced expansion of Kmt2d-deficient cells and altered expression of differentiation markers during acute infection might suggest Kmt2d is required for efficient effector differentiation and survival, similar to what has been reported for CD8 T cells^52^, although to a lower extent in anti-viral responses^53^. In contrast, loss of Kmt2d during contraction/at early memory timepoints might prevent progression to terminal states that are more likely to be deleted. This would conceptually resemble the role of Kmt2d in B cell lymphomagenesis, where a block in differentiation results in prolonged germinal center reactions^54,55^. These data suggest that a loss-of-function mutation may confer a selective advantage or disadvantage on NK cell clones depending on when it is acquired relative to infection. Apart from the specific biology of Kmt2d, which we only assess on the level of population dynamics, our study also provides proof-of-concept that the analysis of somatic mutations can validate the physiological relevance of established targets and identify novel regulators of NK cell fitness. Intriguingly, *KMT2D* disruption was independently discovered to enhance NK cell in vitro persistence at low doses of IL-15 in a recently published genome-wide CRISPR screen^56^, suggesting combined ex vivo genetic and in vitro functional screening approaches could become powerful tools for the discovery of novel regulators of NK cell fitness.

Our reconstruction of clonal age and expansion dynamics further raises important questions on the precise dynamics of clonal selection, as well as the mechanisms for long-term clonal maintenance. The predicted growth period spanning several months to years is an intriguing parallel to “inflationary” CMV-specific T cell responses^57^. In vivo studies have shown that these can be characterized by successive clonal sweeps in the first months after infection^58^. Once established, the half-life and self-renewal of CMV-specific memory T cells are similar to other specificities when taking differentiation state into account, suggesting most dynamics occur early on, followed by relatively stable maintenance^59^. Our mutational-clock based predictions of clonal age intriguingly coincide with CMV-seroconversion rates, which might point towards similar dynamics for memory NK cells. Longitudinal studies during and following acute CMV infection will be instrumental to resolve this question.

Overall, these findings underline the remarkable impact of clonal selection on population dynamics of innate lymphocytes. Future studies will need to establish whether clonal selection is the rule rather than the exception also for NK cells in different settings than CMV infection, where the unique environment and antigenic kinetics expand clones with characteristic phenotypes to sizes detectable with current approaches. Such stringent selection of immune cells with invariant receptors seems like a plausible requirement for the evolution of directed genetic recombination systems that have at least emerged twice in vertebrates^60^, and suggests that clonal selection in the innate immune system might be more widespread than currently appreciated.

## Methods

### Ethics statement and sample collection

Samples were either collected from healthy donors at Charité - Universititätsmedizin Berlin or acquired as buffy coats from Deutsches Rotes Kreuz Nord-Ost. The use of these samples was approved by the Charité ethics commission (EA1/243/24, EA4/270/21 EA1/057/25).

### HCMV serology

For buffy coats, HCMV serology was performed at DRK Dresden. Serological status of fresh blood donors was analyzed by HCMV IgG enzyme-linked immunosorbent assay (IBL International) following the manufacturer’s instructions.

### Sample Processing

Peripheral blood mononuclear cells were isolated by density gradient centrifugation (Ficoll Paque Plus, GE Healthcare) and were first either depleted of T cells using CD3 Microbeads (Miltenyi Biotec) or cryopreserved as total fraction in FBS containing 10% DMSO.

### Flow cytometry

Cell suspensions were stained with combinations of the following fluorochrome-conjugated anti-human antibodies: CD56 PE/Dazzle 594 (1:200), CD3 PE-Cy5 (1:200), CD3 NIR685 (1:100), CD2 PerCp-Cy5.5 (1:200), KLRG1 KIRAVIA Blue 520 (1:200), CD7 BV711 (1:100), NKG2A BV605 (1:50), CD14 BV510 (1:200), CD19 BV510 (1:200), and CD57 BV421 (1:200) (all BioLegend); CD7 BV786 (1:25), CD56 BUV805 (1:800), CD3 BUV805 (1:50), CD16 BUV496 (1:50), and KIR3DL1 BV650 (1:100) (all BD Biosciences); CD159a (NKG2A) biotin/PE-Vio770 (both 1:50) and CD159c (NKG2C) PE (1:100), NKp30 PE-Vio770 (1:50), KIR2DL2/L3 PE-Vio615 (1:25), Siglec-7 APC-Vio770 (1:25) (all REAfinity, Miltenyi Biotec); CD3 APC-eFluor 780 (1:50), CD14 APC-eFluor 780 (1:50), and CD19 APC-eFluor 780 (1:50) (all Thermo Fisher); ILT2 A647 (1:50), CD161 SBUV700 (1:25) (both BioRad); or anti-mouse antibodies: CD49b PE-Dazzle594 (1:200), CD45.1 PacificBlue (1:400), CD45.2 A700 (1:400), F4/80 APC-Cy7 (1:400), (all BioLegend), NK1.1 APC (1:100), CD3 APC-Cy7 (1:200), Ly6G APC-Cy7 (1:400), (all Tonbo Biosciences), CD11b PerCp-Cy5.5 (1:400), Ly49H FITC (1:200), CD19 APC-Cy7 (1:800), CD27 PE-Cy7 (1:1000) (all eBioscience). Dead cells were excluded using Fixable Viability Dye eFluor780, eFluor506 (Thermo Fisher) or Zombie Aqua Fixable Viability kit (BioLegend). Stainings were performed in phosphate-buffered saline (PBS), washed and resuspended in PBS with 1% BSA. Data were acquired on an LSR Fortessa, Symphony A2 (both BD Biosciences), and ID7000E (Sony Biotechnology). FlowJo v10 was used for analysis of flow cytometry data, and GraphPad Prism v10.6.1 was used for statistical analysis.

### DNA extraction from sorted NK cells and control populations

Cryopreserved PBMCs were thawed and stained with the following fluorochrome-conjugated antibodies: CD7 BV711 (1:100), CD3 BV605 (1:50), CD57 BV421 (1:50), CD14 PerCp-Cy5.5 (1:400), and CD19 BV510 (1:200) (all BioLegend); and CD159a (NKG2A) VioBright B515 (1:50) and CD159c (NKG2C) PE (1:100) (both REAfinity, Miltenyi Biotec). Monocytes were sorted as CD14^+^, total NK cells as CD7^+^ CD3^-^ CD14^-^ CD19^-^ or enriched as NKG2C^+^ NKG2A^-^ “memory” and NKG2A^+^ NKG2C^-^ CD57^-^ “conventional” subsets using MA900 (Sony Biotechnology), FACSAria II (BD Biosciences), and Aurora CS (Cytek) cell sorters. Cells were sorted into PBS/BSA and frozen as cell pellets for DNA extraction using silica columns (Qiagen QIAamp DNA Micro) following the manufacturer’s instructions. Alternatively, cells were sorted directly into lysis buffer (30 mM Tris-HCl pH 8.0, 0.5% Tween-20, 0.5% IGEPAL CA-630), adapted from a published protocol^61^. These were digested with Proteinase K (1:20, Qiagen) and Monarch RNase A (1:200, NEB) at 56 °C for 15 minutes, followed by extraction of genomic DNA using SPRIselect or Ampure beads. Buccal swabs were collected using sterile OmniSwabs (Qiagen) and extracted using silica columns (Qiagen QIAamp DNA Blood Mini) following the manufacturer’s instructions.

### Exome library preparation and target capture

Whole-exome sequencing was performed by commercial service providers (CeGaT Tübingen or Novogene Europe). Target enrichment used the Twist Exome 2.0 kit (CeGaT) for 71 donors and the Agilent SureSelect Human All Exon v6 kit (Novogene) for the remaining 10 donors. Libraries were multiplexed and sequenced on Illumina NovaSeq 6000 instruments, 2 x 100 bp for Twist and 2 x 150 bp for Agilent.

### Exome library read processing and alignment

All cohorts were processed with a single Nextflow pipeline (imprint; https://github.com/ollieeknight/imprint). Adapters were trimmed with fastp^62^ v1.3.5 (--detect_adapter_for_pe, --length_required 36). Surviving reads were aligned to the GRCh38 no-alt analysis set with bwa-mem3^63^ v0.6.0 (-Y), merged and coordinate-sorted with samtools^64^ v1.23.1, and duplicate-marked with GATK^65^ v4.6.2.0 MarkDuplicates. Overlapping mate segments were reconciled with fgbio v4.1.0 CallOverlappingConsensusBases, retaining the higher-quality base at disagreements so each fragment contributed once per site.

### Somatic variant calling

Each NK subset was called against its donor’s monocyte control (capture targets padded ±50 bp), with three callers in parallel: (i) GATK Mutect2, with the gnomAD v4.1.1 germline resource and the GATK 1000 Genomes panel of normals (--pcr-indel-model AGGRESSIVE --max-mnp-distance 2). Soft-clipped bases were excluded only when the fastp insert-size peak was at or below the mean read length minus 10 bp. Calls were filtered with FilterMutectCalls using LearnReadOrientationModel and GetPileupSummaries/ CalculateContamination. (ii) Manta^66^ v1.6.0 and Strelka2^67^ v2.9.10 in --exome mode, with Manta’s candidate indels passed to Strelka2 as breakpoint priors. (iii) DeepSomatic^68^ v1.10.0, whole-exome model.

The union of PASS calls was retained. Records were left-aligned and multi-allelic/MNP sites decomposed with bcftools^64^ v1.23.1 norm, each split record tagged with its parent for cross-caller comparison. Allele fractions and depths were recomputed at every union site with vafator^69^ v3.0.0 (MAPQ/BQ≥20), excluding ambiguous bases.

### Annotation and filtering

Ensembl VEP^70^ v116 (--canonical --mane --pick) was used with one consequence per variant. AlphaMissense^32^, SpliceAI^71^ and dbNSFP^72^ v5.31a (REVEL) supplied missense/splicing predictions; NMD, SpliceRegion and pLI supplied nonsense-mediated decay, splice-region and loss-of-function intolerance annotations. Population frequencies were assigned from gnomAD v4.1.1 and somatic identifiers from COSMIC v104.

Alignment properties were estimated per sample, and observations were extracted for both members of the pair. Union sites were classified with Varlociraptor^73^ v8.9.5 under a paired tumor/normal scenario (heterozygosity 1×10^-3^, somatic mutation rate 1×10^-6^ in both samples; sex chromosome ploidy from somalier-inferred sex). Kit-matched panels of normals were built from the monocyte libraries (Mutect2 in tumor-only mode with CreateSomaticPanelOfNormals, --min-sample-count 2). Candidate variants required: absence from the kit-matched panel of normals, ≥3 supporting alt reads, gnomAD v4.1.1 AF ≤ 1×10^-4^, no read-position bias (vafator rank-sum *P* ≥ 0.05) and indels ≤ 10 bp. Surviving candidates passed 1% local FDR control (1 − P(absent) − P(artifact) > 0.99 and somatic posterior > 0.5), or were called by more than one caller. Variants duplicated within cohorts across donors were removed as likely artifacts.

Variants included in mutation load and clonal size analysis were further filtered to be within unpadded capture targets, with VAF distributions visualized for all variants exceeding a threshold of 3% to include the end of the polyclonal tail while optimally resolving clonal peaks visually.

### Selection analysis

Selection was quantified using dndscv^30^ v0.0.1.0 with the provided GRCh38 reference CDS and covariates.Following the authors’ guidance on artifacts, mutations at contiguous positions within a donor were removed before fitting. Global dN/dS ratios were computed for missense, nonsense, splice-site, truncating and all substitution classes, with 95% confidence intervals and likelihood-ratio P values. Ratios for COSMIC Cancer Gene Census v101 tier 1 genes were fitted with genesetdnds, which models the gene set and the remaining genes as separate arms.

### HLA and KIR genotyping

Donor HLA class I genotypes were called from merged donor BAMs with OptiType (--dna), and KIR genes genotyped and copy number estimated using kir-mapper. Self-specific KIR-ligand status per donor was assigned from the OptiType HLA-C genotype (C1/C2 grouping) and HLA-B genotype (Bw4 status) combined with the donor’s kir-mapper KIR gene calls.

### Single cell mutation profiling

Single-cell DNA sequencing was performed using the Mission Bio Tapestri platform according to the manufacturer’s instructions. Custom panels were designed for each individual by performing additional WES on the total CD56^dim^ NK cell compartment and combining all mutations identified from WES of memory, conventional, and total CD56^dim^ populations. As mutation numbers only slightly exceeded 100 variants, we removed surplus mutations with the lowest VAFs until 100 mutations per donor successfully passed the Tapestri Designer Pipeline.

For simultaneous detection of proteins and mutations, cells were stained with the following fluorochrome- and TotalSeq-D oligonucleotide-conjugated antibodies in Cell Staining Buffer (BioLegend) including Human TruStain FcX (BioLegend) and Blocking Buffer (MissionBio): CD7 BV786 (1:25, BD Biosciences), CD19 APC-eF780, CD14 APC-eF780 (eBiosciences, 1:50), CD3 BV605 (BioLegend, 1:100); TotalSeq-D0149 CD161, D0168 CD57 (1:50); D0047 CD56, D0153 KLRG1, D0801 NKp30, D5026 KIR2DL2/L3/S2 (1:100); D5025 KIR2DL1/S1/S3/S5, D0367 CD2, D1374 NKG2A, D1045 CD160, D0083 CD16 (1:200); D1375 NKG2C, D5028 KIR3DL1 (1:400). Cells were washed twice and NK cells purified by fluorescence-activated cell sorting (Viable CD3^-^ CD14^-^ CD19^-^ CD7^+^). NK cells were then encapsulated into microfluidic droplets for lysis and barcoding, followed by multiplex PCR amplification of targeted genomic regions and antibody-derived tags. Barcoded amplicons were recovered, purified, and prepared for sequencing according to standard library preparation workflows. Libraries were sequenced on an Illumina NextSeq 2000 (PE150). Quality control parameters were sufficient for all three analyzed individuals: mean reads/cell/amplicon (HC02: 107, HC09: 117, HC12: 245), panel uniformity (HC02: 67%, HC09: 67%, HC12: 67%), reads mapped to target (HC02: 94%, HC09: 96%, HC12: 92%), and mean reads/cell/antibody (HC02: 752, HC09: 386, HC12: 2049).

### Single cell mutation profiling data analysis

Sequencing data were processed using the Tapestri Pipeline (Mission Bio) for read alignment, cell barcode assignment, variant calling, and quantification of antibody-derived tags. Outputs from the Tapestri Pipeline v3.4 were imported and pre-processed using the manufacturer-provided Tapestri Mosaic v3.7. Briefly, variants were initially filtered using minimum depth and genotype-quality thresholds of 10 and 30, respectively, requiring informative calls in at least 50% of cells and mutation detection in at least 0.5% of cells. Variants identified by this filtering or included in predefined whole-exome and mitochondrial whitelists were retained. Variants were annotated using VarSome, protein counts were normalized using the noise-to-signal protein method, and processed DNA and protein matrices were exported for downstream analysis in R.

Using Seurat v5.4.0^74^, per-cell variant allele frequencies, nucleotide genotype calls and normalized protein counts were incorporated into separate DNA, NGT and ADT assays. Per-cell allelic dropout rates were calculated as the proportion of loci with missing variant allele-frequency measurements.

Variants were initially retained when their median genotype quality across cells was at least 40 and their median sequencing depth was at least 10 reads. The resulting variants were restricted to loci identified by bulk whole-exome sequencing. For clone identification, only variants detected in more than 0.5% of cells and with an allelic dropout rate below 5% were retained. Additional technically or biologically non-informative variants were excluded manually.

Genotype calls were binarized according to the presence or absence of a variant, with heterozygous and homozygous alternative genotypes both encoded as mutated. Missing genotype calls were treated as wild-type for this analysis. A two-dimensional UMAP embedding was calculated directly from the binary genotype matrix using Hamming distance, 30 nearest neighbors and a minimum-distance parameter of 0.2. A shared nearest-neighbor graph was constructed from the UMAP coordinates, and cells were clustered using the Louvain algorithm. Cluster-associated variants were identified from the variant allele-frequency matrix using Seurat’s FindAllMarkers function. Clusters lacking defining mutations were merged with the non-clonal population.

To infer relationships among genetically defined subclusters, the mean binary mutation state of each cluster was calculated. A mutation was considered present within a cluster when its mean binary value exceeded 0.8. Pairwise binary distances between clusters were calculated and subjected to hierarchical clustering using Ward’s minimum-variance method. Cluster order was defined according to the resulting dendrogram. Individual subclusters sharing nested mutation patterns were subsequently combined into broader clonal families, while retaining the fine subcluster assignments for visualization of intraclonal structure.

Following final cluster assignment, cluster-associated variants were recalculated using a minimum log-fold-change threshold of 0.05 and requiring detection in at least 20% of cells in the respective cluster. Mutations were ordered according to their distribution across the inferred clonal families and displayed as heatmaps of single-cell variant allele frequencies. Normalized surface-protein measurements were scaled and visualized across the genetically defined clusters.

Mitochondrial mutations were analyzed using a custom implementation of the mgatk package^75^. To process targeted Mission Bio Tapestri data, spatial read deduplication was disabled to preserve true amplicon coverage, with variant identification accuracy improved by strictly parsing CIGAR insertions to prevent alignment-induced coordinate shifts. A depth-aware strand bias filter was also introduced to prevent variant dropout, with outputs filtered for variants confidently identified in paired ASAP-seq data.

### Construction of phylogenetic trees

Clonal phylogenies were reconstructed from single-cell genotyping data generated using the Mission Bio Tapestri platform. Processed data were analyzed in R using Seurat. To resolve subclonal structures for each donor, genotype matrices (NGT0 assay) were aggregated at the cluster level by averaging across cells, followed by binarization (presence/absence of mutations using a threshold of 0.8). Variants and clusters lacking detectable mutations were excluded, yielding a binary mutation matrix representing subclonal genotypes.

Phylogenetic trees were inferred using infSCITE^76^. Resulting tree structures were parsed from graph (.gv) and Newick outputs to extract mutation assignments and sample attachments. Subclonal abundances were derived from cluster frequencies in the original single-cell data and mapped onto the inferred phylogeny. Where multiple subclones mapped to the same phylogenetic node, clone sizes were aggregated.

Phylogenies were visualized using ggtree^77^ v4.0.4. Nodes were annotated with mutation and clone information, and subclone sizes were represented by scaled node markers proportional to their relative abundance.

### ASAP-Seq with mtDNA recovery

NK cells were enriched from peripheral blood mononuclear cells isolated from freshly collected blood by magnetic depletion using microbeads targeting CD3 and CD14 and cryopreserved in FCS with 10% DMSO. After thawing, cells were stained with the following fluorochrome-conjugated antibodies: CD7 BV786 (1:25; BD Biosciences); CD3 APC-eFluor 780 (1:50), CD14 APC-eFluor 780 (1:50), and CD19 APC-eFluor 780 (1:50; all Thermo Fisher); and CD159a (NKG2A) biotin (1:50) and CD159c (NKG2C) PE (1:100; both REAfinity, Miltenyi Biotec). Fc receptors were blocked with Human TruStain FcX (1:50; BioLegend) for 15 min at 4 °C, and dead cells were excluded using Fixable Viability Dye eFluor 780 (Thermo Fisher).

In a second staining step, cells were incubated for 30 min at 4 °C with combinations of the following nucleotide barcode-labeled antibodies (BioLegend): TotalSeq-A0084 CD56 (1:200), A0083 CD16 (1:500), A0436 anti-biotin (1:100), A0147 CD62L (1:100), A0168 CD57 (1:100), A0367 CD2 (1:1,000), A0801 CD337 (NKp30; 1:100), A0149 CD161 (1:100), A0420 CD158 (1:100), A0592 CD158b (1:100), A0599 CD158e1 (1:100), A0902 CD328 (Siglec-7; 1:1,000), A0867 CD94 (1:100), A0896 CD85j (ILT2; 1:100), A0911 anti-phycoerythrin (1:100), A0251 hashtag 1, A0252 hashtag 2, A0253 hashtag 3, A0254 hashtag 4, A0255 hashtag 5, A0256 hashtag 6, A0257 hashtag 7, A0258 hashtag 8, A0259 hashtag 9, A0260 hashtag 10, A0262 hashtag 12, A0263 hashtag 13, A0264 hashtag 14, and A0265 hashtag 15 (all 1:200–1:400).

As described in the ASAP-seq and mitochondrial scATAC-seq protocols^23,75,78^, cells were fixed with 1% paraformaldehyde for 10 min. Fixation was quenched by adding glycine to a final concentration of 0.125 M, followed by two washes with PBS containing bovine serum albumin (BSA). NK cells were sorted on a FACSAria II (BD Biosciences) as viable, single CD3- CD14- CD19- CD7+ cells.

After sorting, cells were lysed for 3 min on ice in modified lysis buffer containing 10 mM Tris-HCl (pH 7.5), 10 mM NaCl, 3 mM MgCl_2_, 0.1% NP-40, and 1% BSA. Lysed cells were subsequently washed with buffer containing 10 mM Tris-HCl (pH 7.5), 10 mM NaCl, 3 mM MgCl_2_, and 1% BSA, and resuspended in diluted nuclei buffer (10x Genomics). Successful lysis was confirmed by trypan blue staining, after which cells were counted and processed as described below.

scATAC-seq libraries were prepared using the Chromium Next GEM Single Cell ATAC Reagent Kit v2 according to the manufacturer’s instructions, with modifications described in the ASAP-seq and mitochondrial scATAC-seq protocols to enable recovery of TotalSeq antibody-derived tags (ADTs), hashtag oligonucleotides (HTOs), and mtDNA. The eluate retained from the silane bead elution was combined with the supernatant from the first SPRIselect purification and used to amplify ADT/HTO libraries with KAPA HiFi ReadyMix (Roche) and sample-specific index primers (Illumina small RNA RPIx/TruSeq D7xx), followed by purification with SPRIselect reagent. Library size and quality were assessed using a Fragment Analyzer (Advanced Analytical) before fragmentation and after final purification. Final library concentrations were measured using a Qubit 2.0 fluorometer (Thermo Fisher). Libraries were sequenced on a NovaSeq 6000 instrument (Illumina) using R1 100 cycles, R2 100 cycles, i7 8 cycles, i5 16 cycles.

### ASAP-Seq data analysis

Analysis of previously published single-cell ATAC-seq data (GSE197037) was performed as described^19^. New data generated here were analyzed in R using Seurat^74^ v5.4.0 and Signac^79^ v1.16.0. Fragment files and the peak set generated by the Cell Ranger ATAC pipeline were used to construct a peak-by-cell count matrix and a Seurat object. Cells with fewer than 500 total fragments or invalid barcodes were excluded during data import. Genotype-based donor assignments were obtained using Vireo^80^, and multiplets identified by AMULET^81^ were removed. Antibody-derived tag (ADT) and hashtag oligonucleotide counts were added as separate assays and normalized using centered log-ratio transformation.

Quality-control metrics included the number of fragments in peaks, transcription start-site enrichment, nucleosome signal and the fraction of reads mapping to ENCODE blacklist regions. Cells were retained when they had more than 1,000 ATAC fragments, a TSS enrichment score greater than 4, a nucleosome signal of 2 or less and a blacklist fraction below 0.002. Gene annotations were obtained from EnsDb.Hsapiens.v86 and converted to the hg38 UCSC chromosome nomenclature.

Chromatin accessibility data were normalized by term frequency–inverse document frequency transformation, followed by singular-value decomposition. Uniform manifold approximation and projection was calculated from latent semantic indexing components 2–30. Cells were clustered using a shared nearest-neighbor graph with the Louvain algorithm. Gene activity scores were calculated with Signac and log-normalized. Non-NK-cell contaminants were identified from ADT expression and gene activity profiles and excluded. Peaks were then recalled in the retained NK cells using MACS3^82^, and donor-specific datasets were reprocessed and clustered separately. Cluster annotations and selected subclusters were refined based on accessibility-derived gene activity and ADT expression of NK-cell differentiation and receptor markers.

Mitochondrial allele counts generated with mgatk^75^ were added to the Seurat objects. Cells with a mitochondrial sequencing depth of 5 or less were excluded. Variable mitochondrial sites were identified using Signac, and high-confidence variants were required to be confidently detected in at least five cells, have a strand correlation of at least 0.65 and exceed the donor-specific variance-to-mean ratio threshold. Per-cell mitochondrial variant allele frequencies were subsequently calculated for the retained variants and used to assess mitochondrial clonal structure as previously described^19^. Briefly, cells were grouped by mitochondrial allelic profiles using FindClonotypes with Euclidean distance, followed by graph-based clustering. Clonotypes were screened for enriched mitochondrial variants using FindAllMarkers. Clonotypes supported by enriched variants were assessed for association with independently defined NK-cell clusters using χ^2^ statistics and an FDR of 0.05. Clonotypes associated with the adaptive NK-cell compartment were selected following inspection of their distribution in the UMAP embedding. For frequency estimation, analyses were restricted to clusters corresponding to the NKG2C^+^ adaptive compartment. Individual clone frequencies were calculated relative to all cells in this compartment, including cells outside the selected clonotypes. The cumulative clonal fraction was calculated as the sum of the selected clone frequencies within each donor.

Mitochondrial subclones within the ASAP-Seq dataset were mapped to scDNA-seq-defined clonal families by using mutations identified across both assays as anchoring points. In addition to subclones that exhibited fully overlapping variants between the datasets, ASAP-seq of donor HC02 also revealed a specific clonotype (defined by *1485G>A*) nested within a larger mitochondrial subclone (*7395T>G*), so we included cells carrying either variant to maximize the number of cells available for the analysis of clone D in the ASAP-seq dataset.

### Whole Genome Sequencing

WGS libraries were prepared by a commercial provider (CeGaT GmbH Tübingen) using the TruSeq DNA Nano library preparation kit (Illumina) and 100 ng genomic DNA as input. Libraries were sequenced on a NovaSeq X Plus (PE150) achieving an average coverage of 90x for the NK cell subsets or 30x for monocyte and buccal swab controls. Demultiplexing was performed with Illumina bcl2fastq (version 2.20).

### Read processing, alignment and mutation calling from WGS data

Raw sequencing reads from NK cells and matched monocyte or buccal swab control samples were 5’ and 3’ trimmed using fastp^62^. Reads were mapped to the human hg38 reference genome using bwa-mem^83^ v0.7.18. Bam files were cleaned with gatk cleansam^65^ v4.6.1.0 and coordinate-sorted using samtools^84^ sort v1.20. Duplicate reads were marked with gatk markduplicates v4.6.1.0 and bam files were indexed using samtools index v1.20.

Somatic SNVs and small insertions/deletions (indels) were called using Strelka v2.9.10 and gatk Mutect2 v4.2.0.0 in tumor-normal mode, with matched buccal swab samples serving as germline control. Variants were filtered by internal gatk Mutect2 filters and applying a read-orientation filter. All variants were annotated with ANNOVAR^85^ according to the human reference genome version hg38. Variants located in repeat regions and simple repeat regions were filtered using bedtools^86^ intersect v2.31.0. Finally, the intersection of Mutect2 and Strelka outputs was computed. VAFs of SNVs and indels were calculated as the number of variant reads divided by the sum of variant reads and reference reads.

### Extraction of single base substitution signatures

Somatic single-nucleotide variants identified by WGS were formatted as SigProfiler input, with samples defined by cell type and donor. Mutational count matrices were generated using SigProfilerMatrixGeneratorR^87^ v1.2.13. Counts were aggregated across chromosomes, and mutational spectrum plots were generated. Mutational signatures were extracted de novo from the SBS96 mutation count matrix using SigProfilerExtractor^88^ v1.1.25. Solutions comprising one to five signatures were evaluated using 100 non-negative matrix factorization replicates per signature number.

### Inference of model parameters from WGS/WES data

We used a new version of SCIFER to infer model parameters from WES and WGS data (manuscript in preparation). To obtain reliable results, SCIFER requires coverage and variant-allele-frequency information of true-positive, disomic variants. From each sample, we thus selected autosomal variants with variant allele frequencies (VAFs) greater-or-equal to 0.05. In SCIFER, we binned the VAFs using the setting Δread = 3 and the OligoCloneDynamics model with a single clone. Parameter inference was done using SCIFER’s ABCSMC function, terminating when either the acceptance rate dropped below 0.001 or 50 generations were simulated.

For the WES data (n = 81), we performed parameter inference only if at least 10 variants were found (n = 76). Furthermore, in order for a group of variants to be recognized as the leading clone, this group must contain at least 3 variants.

### Classification of clones

Parameter inference for the WES data delivered an estimate for the clone size and the number of (exome-wide) founder mutations for each donor (Extended Data Fig. 2a). While the SCIFER model was initialized as a one-clone model, it is in principle conceivable that several clones grew to a similar size. In such a case, SCIFER is not able to infer the actual number of clones, rather a single clone with a large number of founder mutations is inferred. Clearly, this issue cannot arise if the clone size is larger than 50%. The scatter plot shown in Extended Data Fig. 2a shows an L-shape, suggesting that clones of size above 25% are indeed monoclonal (red dots), whereas clones with smaller size (grey dots) are often oligoclonal (although some might also be monoclonal).

### Estimation of age-at-clonal-origin distribution

The age at clonal origin was estimated based on donors with a single dominant clone. To assure monoclonality, we selected all donors from the WES cohort with a dominant clone of clone frequency above 25% (n = 49, Extended Data Fig. 2a). This sub-cohort includes the three donors for which also WGS data were measured. These three donors were used to estimate a scaling factor between the genome-wide and exome-wide numbers of clonal mutations. Specifically, the ratio of the number of clonal mutations in the WGS data and the WES data was individually computed and subsequently averaged, yielding a value of 32, i.e., on average, 1 in 32 mutations falls into the exome. We multiplied the exome-wide number of mutations by this scaling factor to obtain an estimate for the genome-wide number of mutations for each donor in the selected sub-cohort.

To perform real-timing, we used the measured relationships between mutational burden *M*(*t*) and age *t* from Machado et al^9^. Since these data do not include NK cells, we resorted to naive T cells for which the relationship reads *M*(*t*) = 22 * *t* + 164. Inverting this relation allows us to map the genome-wide number of mutations to the time of origin, which we then binned by decade.

### Expected time to clonality under neutral drift

To estimate the time it would take a cell population to establish clonality without selection, we consider a neutral, time-continuous Moran process. Specifically, let the population size be *N* and the division rate per cell *λ*. Conditioning on one of the initial cells to eventually fixate and hence establish clonality, the expected time to do so is given by (*N*-1) / *λ* ^89^.

### Mice

The mice used in this study were housed at Memorial Sloan Kettering Cancer Center in accordance with the guidelines of the Institutional Animal Care and Use Committee. Mice were bred under specific-pathogen-free conditions in 12 hour light–dark cycles at 72 °F with 30–70% humidity. Age- and sex-matched mice were used for each experiment, with both sexes included throughout the study. The following strains were used in this study: Klra8^-/-^ (Ly49H-deficient^90^, received from Silvia M. Vidal), C57BL/6 CD45.1 (Stem.1, received from David T. Scadden), C57BL/6 CD45.1xCD45.2, Rag2-/- x Il2rg-/- mice (Taconic Model No. 4111), Ncr1-Cre (received from Eric Vivier) x Kmt2d-flox (Kmt2dtm1.1Kaig/J^91^, received from Ari Melnick), Vav1-Cre x Kmt2d-flox, UbcCreERT2 (B6.Cg-Ndor1Tg(UBC-cre/ERT2)1Ejb/1J, The Jackson Laboratory) x Kmt2d-flox tdTomato^LSL^. All experiments were performed in accordance with approved institutional protocols.

### Bone marrow chimeras

To generate bone marrow chimeric mice, CD45.1^+^ (C57BL/6J-CD45.1STEM)/CD45.2+ (C57BL/6J) host mice were subjected to 900 cGy irradiation, followed by adoptive transfer of bone marrow cells incubated with anti-NK1.1 antibody (clone PK136) to deplete mature NK cells. To assess competitive reconstitution kinetics, equal numbers of wild-type and Ncr1-Cre Kmt2d-flox bone marrow cells were co-transferred. For MCMV infection studies, the ratio of bone marrow input was adjusted to achieve approximately 1:1 wild-type:Kmt2d-deficient NK cell reconstitution. Competitive reconstitution was quantified by normalizing to the congenically marked non-NK cell populations.

### MCMV Infection

MCMV (Smith strain) was passaged through BALB/c hosts three times. To generate viral stocks, salivary glands were harvested from mice that had been infected three weeks prior and homogenized. Bone marrow chimera mice were administered 5 x 10^4^ PFU of MCMV intraperitoneally. Klra8^-/-^ mice were administered 1 x 10^3^ PFU of MCMV by intraperitoneal injection.

### Adoptive transfer model

Congenically distinguishable CD45.1- and CD45.2-expressing splenocytes from conditional knockout and wild-type control mice were mixed at approximately equal ratios and co-transferred intravenously into *Klra8^-/-^* recipient mice. One day after transfer, recipient mice were infected with MCMV. The relative expansion of knockout and wild-type NK cells was monitored by weekly peripheral blood sampling. Genotype frequencies at each time point were normalized to their respective frequencies in the transferred input population. For the inducible model using UbcCre-ERT2, 4 mg Tamoxifen in 200 µl corn oil was administered at day 5/7 by oral gavage. Successful induction was monitored by assessing Cre-inducible tdTomato expression.

### Mouse tissue processing and flow cytometry staining

Blood was collected from mice into 100 USP units/mL heparin sodium (BD Biosciences) to prevent coagulation. Spleen tissues were harvested and dissociated using glass slides in staining media (D-PBS, 2% FCS). For bone marrow isolation, one femur and tibia were flushed using a 27-gauge syringe filled with staining media. Samples were treated with ACK lysis buffer (0.15 M NH_4_Cl, 0.01 M KHCO_3_, 0.1 mM NaEDTA) for 5 minutes at room temperature to lyse red blood cells, with blood samples undergoing two rounds of lysis.

### Use of artificial intelligence

Generative AI tools (ChatGPT (OpenAI), Claude (Anthropic)) were used for copy editing of human-generated text to improve clarity, grammar and style. No AI tools were used to generate or analyze data or to draw scientific conclusions. All edits were reviewed and approved by the authors.

## Data Availability

Raw FASTQ files for scATAC and single cell mutations will be accessible through the German Human Genome-Phenome Archive (GHGA; [accession placeholder]) upon publication. Unfiltered mutation calls will be deposited as VCF files. Access to these controlled data requires the submission of a formal project application and subsequent approval by the designated Data Access Committee (DAC). Processed, non-identifiable data will be accessible on Zenodo [accession placeholder].

## Code Availability

Novel custom code used in this study will be available at GitHub [link placeholder] upon publication.

## Acknowledgements

We thank Carolin Hobe, Marion Klemm, and Jennifer Zhang for technical support, and Caleb Lareau and Konstantin Helmsauer for critical discussions.

## Funding

This work was supported by the European Union through the European Research Council Advanced Grant ‘MEM-CLONK’ (101055157 to C.R.). Funding was also provided by the Deutsche Forschungsgemeinschaft under Germany’s Excellence Strategy (EXC 3118/1-533770413), the Priority Programme SPP 1937 (RO3565/4-2), Transregio TRR 241/2-375876048 (project B02), Transregio TRR 412/1 2025-535081457 (project B06), and individual grant RO 3565/7-1 (all to C.R.) and TRR186/A21 (to T.H.). Additional support was provided by the Leibniz-ScienceCampus Chronic Inflammation, the Leibniz-Kooperative Exzellenz (K259/2019 to C.R.), and the Else Kroener-Promotionskolleg Berlin (2024_EKPK.22 to C.R.). Furthermore, this research was funded by the Federal Ministry of Education and Research and the State of Berlin as part of the Excellence Strategy of the Federal Government and the States through the Berlin University Alliance (BUA). This work was further supported by the German Federal Ministry of Research, Technology and Space (BMFTR) (01BIHTP2520B to T.R.) and a Berlin Institute of Health (BIH) SPARK grant (SelectNK to T.R.). T.R. is recipient of an EMBO Scientific Exchange Grant and DAAD short-term fellowship. J.C.S. was supported by the Ludwig Center for Cancer Immunotherapy, the American Cancer Society, the Burroughs Wellcome Fund, and the NIH (AI100874, AI130043, AI155558, AI189793, and P30CA008748). S.G. is supported by the NIAID (NIH) under award number K99AI180360. Views and opinions expressed are those of the authors only and do not necessarily reflect those of the European Union or the European Research Council Executive Agency. Neither the European Union nor the granting authority can be held responsible for them.

## Author contributions

C.R. and T.R. conceived the study. T.R., M.M. and K.J. performed the experiments. M.G. performed population-genetic modelling. T.R., M.G., O.K. and M.M. analyzed the data. T.R., M.G., O.K., T.H. and C.R. wrote the manuscript. S.G. and J.C.S. contributed to study design, data interpretation and supervision of the mouse studies. C.R. and T.H. jointly supervised the study. All authors reviewed and approved the final version.

## Extended Data

**Extended Data Fig. 1.**
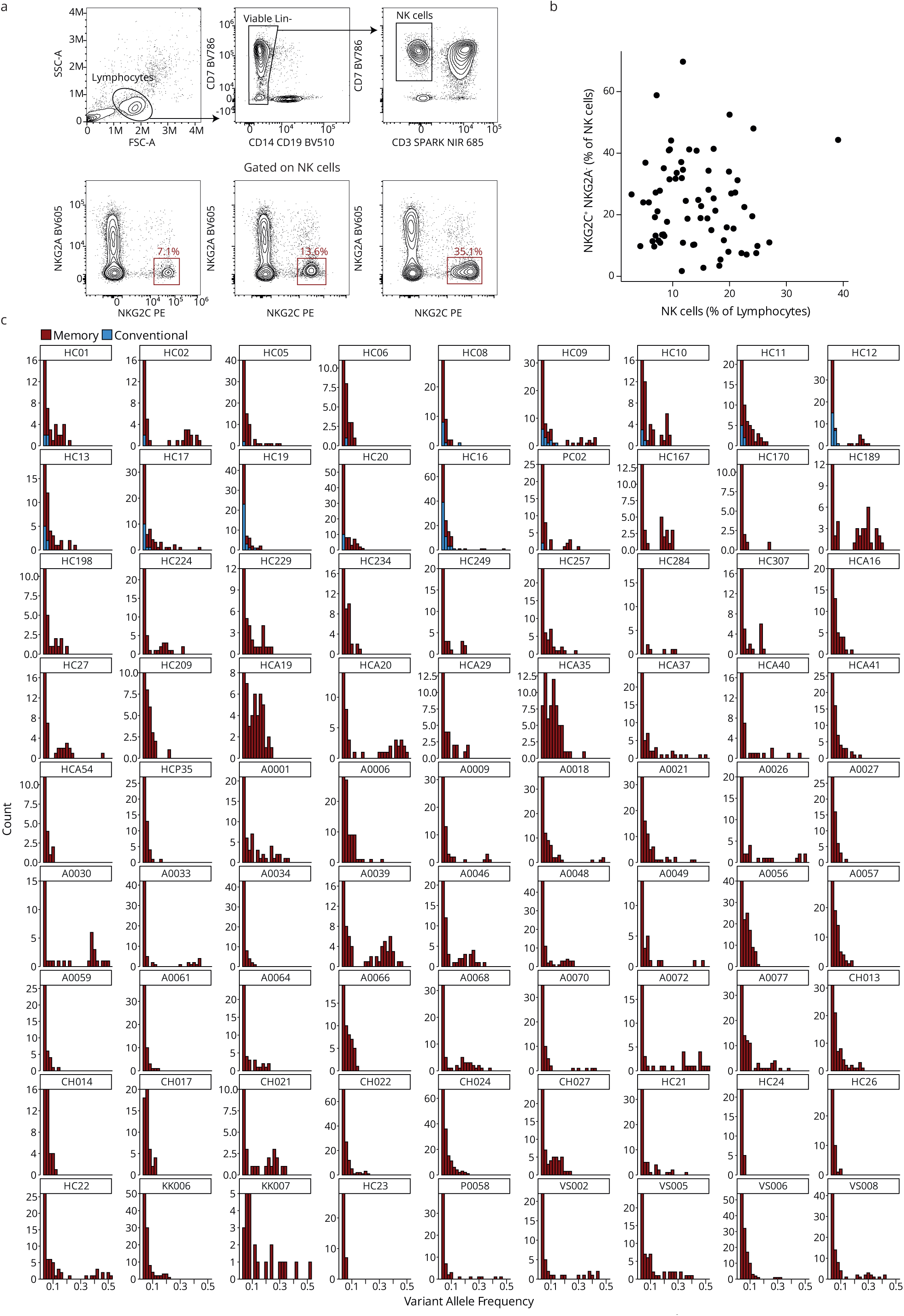
Dominant memory NK-cell clones are widespread in HCMV-seropositive individuals. **a-b** Representatives (**a**) and quantification (**b**) of NKG2C^+^ NKG2A^-^ memory NK cell frequencies in our cohort of HCMV-seropositive individuals (N = 81). **c** Variant allele frequency distributions of memory (NKG2C^+^ NKG2A^-^) and conventional (NKG2A^+^ NKG2C^-^ CD57^-^) NK cell mutations with VAF > 0.03 across different individuals.

**Extended Data Fig. 2.**
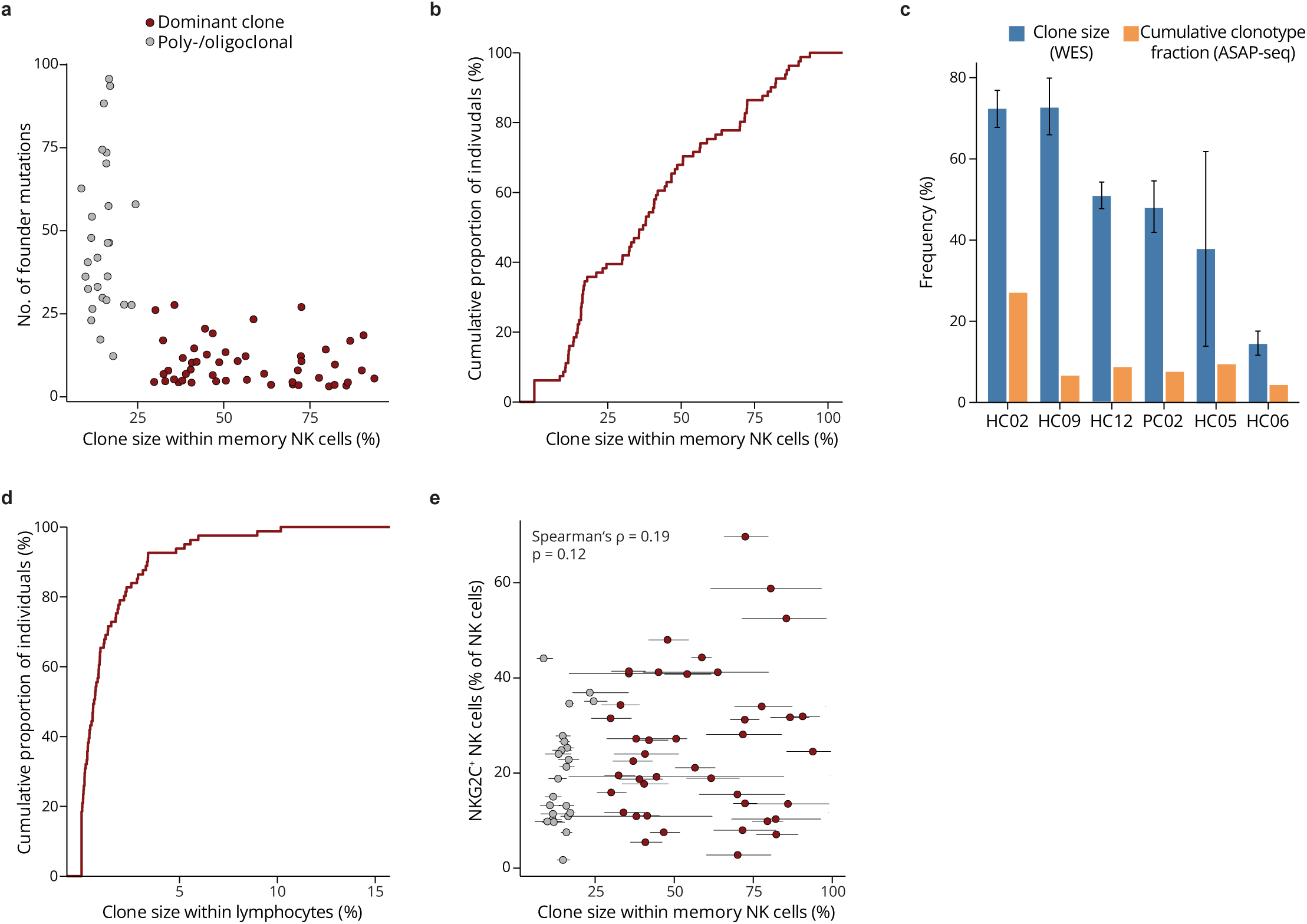
Dominant memory NK-cell clones are widespread in HCMV-seropositive individuals. **a** Estimated number of founder mutations and clone size within memory NK cells from SCIFER^7^. **b** Cumulative size distribution of clones within the memory NK cell compartment (N = 76). **c** Comparison of clone size estimates from WES (with 95% confidence intervals) and cumulative clonotype fraction summing up all memory NK cell clonotypes defined by mitochondrial mutations in ASAP-seq. **d** Cumulative size distribution of memory NK cell clones within lymphocytes (N = 66). **e** Comparison of clone sizes to percentage of NKG2C^+^ cells within total NK cells (N = 66). Statistics from Spearman’s rank correlation.

**Extended Data Fig. 3.**
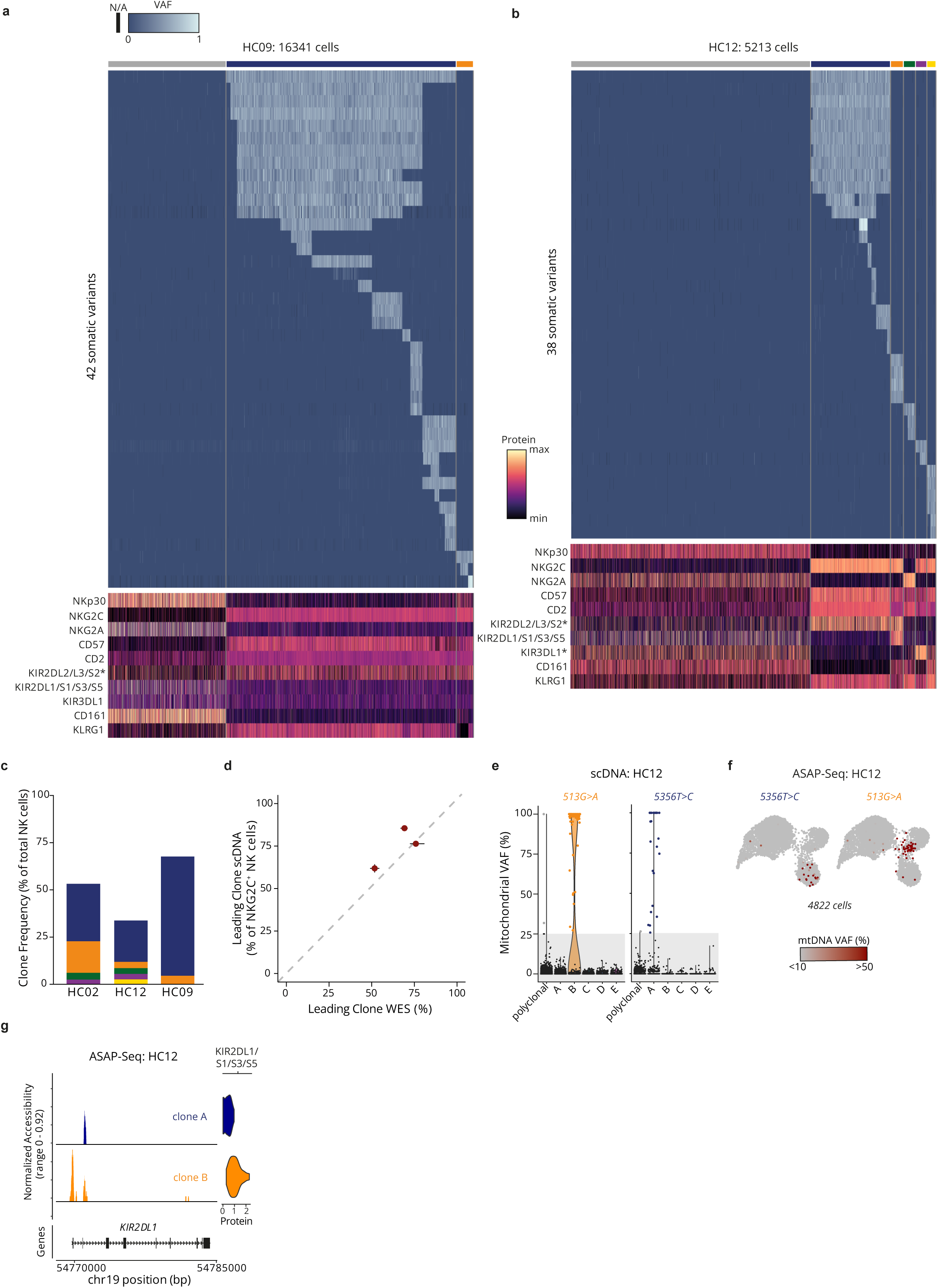
Extensive branching in clonal phylogenies links expansion to stable NK-cell states. **a-b** Joint single-cell profiling of cell surface antigens and somatic mutations in donors HC09 (**a**) and HC12 (**b**). Asterisks mark self-MHC-specific KIRs. **c** Proportions of individual clones within total NK cells **d** Leading clone frequency within NKG2C^+^ NK cells as detected by SCIFER from bulk WES with 95% confidence intervals or targeted single cell mutational profiling (scDNA). **e-f** Clone-associated mtDNA variants in scDNA- (**e**) and ASAP-seq (**f)** for donor HC12. **g** Clone-specific chromatin accessibility profiles at the clonally segregated *KIR2DL1* locus, shown alongside surface protein levels quantified by ASAPseq.

**Extended Data Fig. 4.**
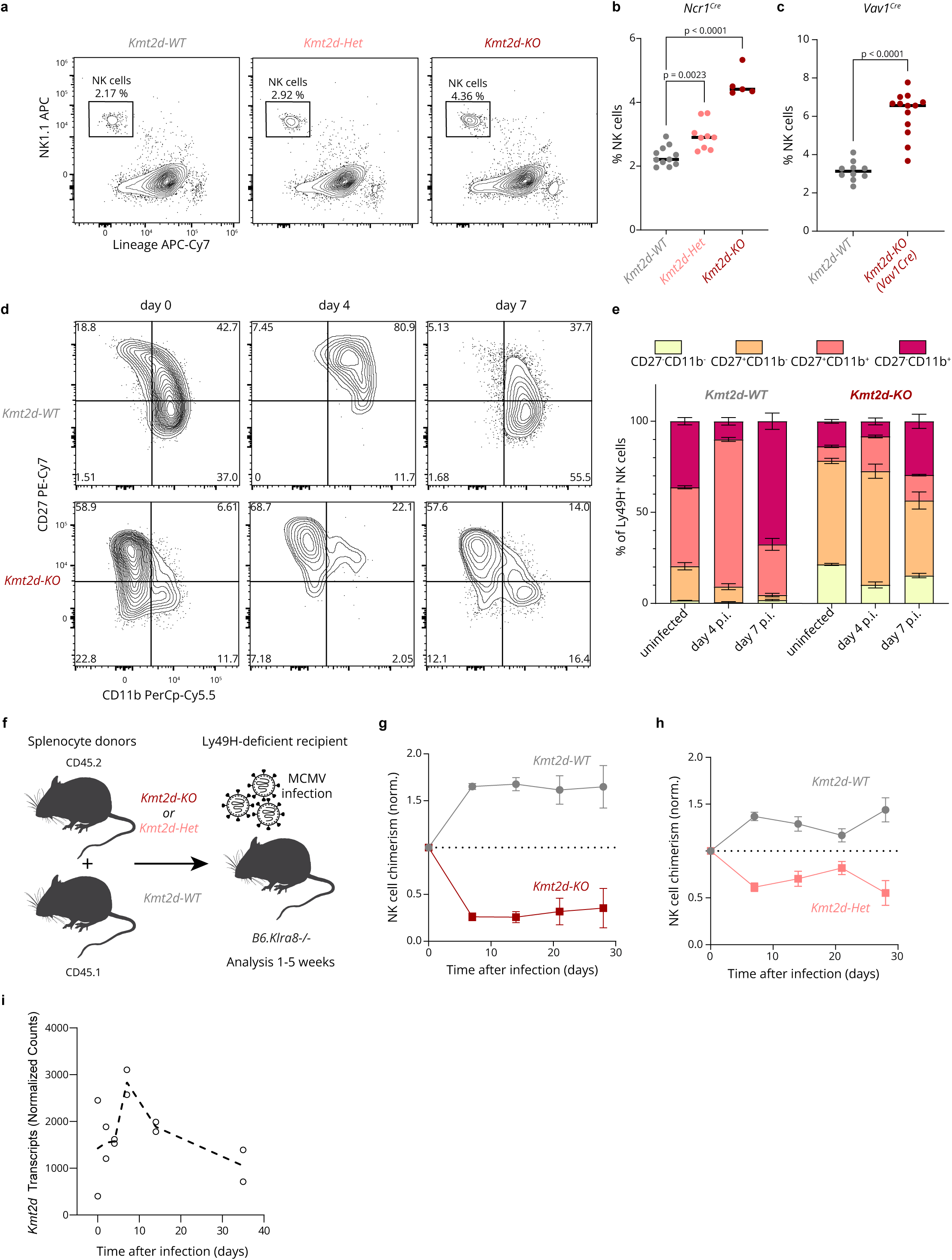
*Kmt2d* loss-of-function is subject to context-dependent selection in vivo. **a-c** Representative NK cell frequencies (a) and quantification (b-c) in spleens of *Ncr1^Cre^*x *Kmt2d^fl/fl^*(Kmt2d-KO), *Ncr1^Cre^*x *Kmt2d^fl/wt^*(Kmt2d-Het), and *Vav1^Cre^*x *Kmt2d^fl/fl^*compared to wild-type (WT) littermates at steady-state. Statistical significance from ordinary one-way ANOVA with Dunnett’s post-test. **d-e** Representative stainings (d) and quantification (e) of CD27 and CD11b on NK cells from mixed bone marrow chimeras at different time points before and after MCMV infection (N=5-8). **f** Congenically marked *Ncr1^Cre^* x *Kmt2d^fl/fl^* (Kmt2d-KO) or *Ncr1^Cre^* x *Kmt2d^fl/wt^* (Kmt2d-Het) splenocytes were co-transferred with *Ncr1^Cre^*x *Kmt2d^wt/wt^*(WT) into Klra8^-/-^ (encoding Ly49H) hosts before infection with MCMV. **g-h** Relative chimerism of Kmt2d-KO (N=5-11) (g) and Kmt2d-Het (N=5) (h) NK cells after infection with MCMV in the adoptive transfer model. **i** Normalized *Kmt2d* expression from publicly available^43^ RNA-sequencing data over the course of MCMV infection.

## Notes

### Competing Interest Statement

The authors have declared no competing interest.

## References

1. Marzo, A. L. et al. Initial T cell frequency dictates memory CD8+ T cell lineage commitment. Nat Immunol 6, 793–799 (2005).

2. Busch, D. H. & Pamer, E. G. T cell affinity maturation by selective expansion during infection. J Exp Med 189, 701–710 (1999).

3. Buchholz, V. R. et al. Disparate individual fates compose robust CD8+ T cell immunity. Science 340, 630–635 (2013).

4. Gerlach, C. et al. Heterogeneous differentiation patterns of individual CD8+ T cells. Science 340, 635–639 (2013).

5. Coorens, T. H. H. et al. Extensive phylogenies of human development inferred from somatic mutations. Nature 597, 387–392 (2021).

6. Spencer Chapman, M., et al. Lineage tracing of human development through somatic mutations. Nature 595, 85–90 (2021).

7. Körber, V. et al. Detecting and quantifying clonal selection in somatic stem cells. Nat Genet 57, 1718–1729 (2025).

8. Niroula, A. et al. Distinction of lymphoid and myeloid clonal hematopoiesis. Nat Med 27, 1921–1927 (2021).

9. Machado, H. E. et al. Diverse mutational landscapes in human lymphocytes. Nature 608, 724–732 (2022).

10. Komatsu, H., Sierro, S., V Cuero, A. & Klenerman, P. Population analysis of antiviral T cell responses using MHC class I-peptide tetramers. Clin Exp Immunol 134, 9–12 (2003).

11. Karrer, U. et al. Memory inflation: continuous accumulation of antiviral CD8+ T cells over time. J Immunol 170, 2022–2029 (2003).

12. Sylwester, A. W. et al. Broadly targeted human cytomegalovirus-specific CD4+ and CD8+ T cells dominate the memory compartments of exposed subjects. J Exp Med 202, 673–685 (2005).

13. Sun, J. C., Beilke, J. N. & Lanier, L. L. Adaptive immune features of natural killer cells. Nature 457, 557–561 (2009).

14. Béziat, V. et al. NK cell responses to cytomegalovirus infection lead to stable imprints in the human KIR repertoire and involve activating KIRs. Blood 121, 2678–2688 (2013).

15. Schlums, H. et al. Cytomegalovirus Infection Drives Adaptive Epigenetic Diversification of NK Cells with Altered Signaling and Effector Function. Immunity 42, 443–456 (2015).

16. Hammer, Q. et al. Peptide-specific recognition of human cytomegalovirus strains controls adaptive natural killer cells. Nat Immunol 19, 453–463 (2018).

17. Gumá, M. et al. Imprint of human cytomegalovirus infection on the NK cell receptor repertoire. Blood 104, 3664–3671 (2004).

18. Grassmann, S. et al. Distinct Surface Expression of Activating Receptor Ly49H Drives Differential Expansion of NK Cell Clones upon Murine Cytomegalovirus Infection. Immunity 50, 1391–1400.e4 (2019).

19. Rückert, T., Lareau, C. A., Mashreghi, M.-F., Ludwig, L. S. & Romagnani, C. Clonal expansion and epigenetic inheritance of long-lasting NK cell memory. Nat Immunol 23, 1551–1563 (2022).

20. Mujal, A. M., Delconte, R. B. & Sun, J. C. Natural Killer Cells: From Innate to Adaptive Features. Annu. Rev. Immunol. 39, 417–447 (2021).

21. Rebuffet, L. et al. High-dimensional single-cell analysis of human natural killer cell heterogeneity. Nat Immunol 25, 1474–1488 (2024).

22. Apoil, P. A. et al. Reference values for T, B and NK human lymphocyte subpopulations in adults. Data in Brief 12, 400–404 (2017).

23. Mimitou, E. P. et al. Scalable, multimodal profiling of chromatin accessibility, gene expression and protein levels in single cells. Nat Biotechnol 39, 1246–1258 (2021).

24. Williams, M. J., Werner, B., Barnes, C. P., Graham, T. A. & Sottoriva, A. Identification of neutral tumor evolution across cancer types. Nat Genet 48, 238–244 (2016).

25. Alexandrov, L. B. et al. Clock-like mutational processes in human somatic cells. Nat Genet 47, 1402–1407 (2015).

26. Hoehl, S., Berger, A., Ciesek, S. & Rabenau, H. F. Thirty years of CMV seroprevalence-a longitudinal analysis in a German university hospital. Eur J Clin Microbiol Infect Dis 39, 1095–1102 (2020).

27. Lopez-Vergès, S. et al. Expansion of a unique CD57^+^NKG2Chi natural killer cell subset during acute human cytomegalovirus infection. Proc Natl Acad Sci U S A 108, 14725–14732 (2011).

28. Muntasell, A., et al. *NKG2C* zygosity influences CD94/NKG2C receptor function and the NK-cell compartment redistribution in response to human cytomegalovirus. Eur J Immunol 43, 3268–3278 (2013).

29. Liu, L. L. et al. Critical Role of CD2 Co-stimulation in Adaptive Natural Killer Cell Responses Revealed in NKG2C-Deficient Humans. Cell Rep 15, 1088–1099 (2016).

30. Martincorena, I. et al. Universal Patterns of Selection in Cancer and Somatic Tissues. Cell 171, 1029–1041.e21 (2017).

31. Sondka, Z. et al. The COSMIC Cancer Gene Census: describing genetic dysfunction across all human cancers. Nat Rev Cancer 18, 696–705 (2018).

32. Cheng, J. et al. Accurate proteome-wide missense variant effect prediction with AlphaMissense. Science 381, eadg7492 (2023).

33. Pölönen, P. et al. The genomic basis of childhood T-lineage acute lymphoblastic leukaemia. Nature 632, 1082–1091 (2024).

34. Gutierrez, A. et al. Inactivation of LEF1 in T-cell acute lymphoblastic leukemia. Blood 115, 2845–2851 (2010).

35. Kato, M. et al. Frequent inactivation of A20 in B-cell lymphomas. Nature 459, 712–716 (2009).

36. Compagno, M. et al. Mutations of multiple genes cause deregulation of NF-κB in diffuse large B-cell lymphoma. Nature 459, 717–721 (2009).

37. Küçük, C. et al. PRDM1 is a tumor suppressor gene in natural killer cell malignancies. Proc. Natl. Acad. Sci. U.S.A. 108, 20119–20124 (2011).

38. Koskela, H. L. M. et al. Somatic STAT3 mutations in large granular lymphocytic leukemia. N Engl J Med 366, 1905–1913 (2012).

39. Lindeboom, R. G. H., Vermeulen, M., Lehner, B. & Supek, F. The impact of nonsense-mediated mRNA decay on genetic disease, gene editing and cancer immunotherapy. Nat Genet 51, 1645–1651 (2019).

40. Morin, R. D. et al. Frequent mutation of histone-modifying genes in non-Hodgkin lymphoma. Nature 476, 298–303 (2011).

41. Pasqualucci, L. et al. Analysis of the coding genome of diffuse large B-cell lymphoma. Nat Genet 43, 830–837 (2011).

42. Gao, L.-M. et al. Somatic mutations in KMT2D and TET2 associated with worse prognosis in Epstein-Barr virus-associated T or natural killer-cell lymphoproliferative disorders. Cancer Biol Ther 20, 1319–1327 (2019).

43. Lau, C. M. et al. Epigenetic control of innate and adaptive immune memory. Nat Immunol 19, 963–972 (2018).

44. Soerens, A. G. et al. Functional T cells are capable of supernumerary cell division and longevity. Nature 614, 762–766 (2023).

45. Rückert, T. & Romagnani, C. Extrinsic and intrinsic drivers of natural killer cell clonality. Immunol Rev 323, 80–106 (2024).

46. Adams, N. M. et al. Cytomegalovirus Infection Drives Avidity Selection of Natural Killer Cells. Immunity 50, 1381–1390.e5 (2019).

47. Kaiser, B. K., Pizarro, J. C., Kerns, J. & Strong, R. K. Structural basis for NKG2A/CD94 recognition of HLA-E. Proc. Natl. Acad. Sci. U.S.A. 105, 6696–6701 (2008).

48. Anfossi, N. et al. Human NK Cell Education by Inhibitory Receptors for MHC Class I. Immunity 25, 331–342 (2006).

49. Felices, M. et al. Functional NK cell repertoires are maintained through IL-2Rα and Fas ligand. J Immunol 192, 3889–3897 (2014).

50. Mold, J. E. et al. Clonally heritable gene expression imparts a layer of diversity within cell types. Cell Syst 15, 149–165.e10 (2024).

51. Xie, M. et al. Age-related mutations associated with clonal hematopoietic expansion and malignancies. Nat Med 20, 1472–1478 (2014).

52. Kim, J. et al. Lysine methyltransferase Kmt2d regulates naive CD8+ T cell activation-induced survival. Front Immunol 13, 1095140 (2022).

53. Cohen, J. A. et al. KMT2D coordinates antiviral CD4+ T cell responses through opposing effects on T follicular helper and cytotoxic gene expression. Cell Rep 44, 115775 (2025).

54. Ortega-Molina, A. et al. The histone lysine methyltransferase KMT2D sustains a gene expression program that represses B cell lymphoma development. Nat Med 21, 1199–1208 (2015).

55. Zhang, J. et al. Disruption of KMT2D perturbs germinal center B cell development and promotes lymphomagenesis. Nat Med 21, 1190–1198 (2015).

56. Nikolic, I. et al. Enhancing anti-tumor immunity of natural killer cells through targeting IL-15R signaling. Cancer Cell 43, 2034–2050.e11 (2025).

57. Klenerman, P. & Oxenius, A. T cell responses to cytomegalovirus. Nat Rev Immunol 16, 367–377 (2016).

58. Schober, K. et al. Reverse TCR repertoire evolution toward dominant low-affinity clones during chronic CMV infection. Nat Immunol 21, 434–441 (2020).

59. Van Den Berg, S. P. H., et al. Quantification of T-cell dynamics during latent cytomegalovirus infection in humans. PLoS Pathog 17, e1010152 (2021).

60. Pancer, Z. et al. Somatic diversification of variable lymphocyte receptors in the agnathan sea lamprey. Nature 430, 174–180 (2004).

61. Ellis, P. et al. Reliable detection of somatic mutations in solid tissues by laser-capture microdissection and low-input DNA sequencing. Nat Protoc 16, 841–871 (2021).

62. Chen, S., Zhou, Y., Chen, Y. & Gu, J. fastp: an ultra-fast all-in-one FASTQ preprocessor. Bioinformatics 34, i884–i890 (2018).

63. Vasimuddin, Md., Misra, S., Li, H. & Aluru, S. Efficient Architecture-Aware Acceleration of BWA-MEM for Multicore Systems. in 2019 IEEE International Parallel and Distributed Processing Symposium (IPDPS) 314–324 (IEEE, Rio de Janeiro, Brazil, 2019). doi:10.1109/IPDPS.2019.00041.

64. Danecek, P. et al. Twelve years of SAMtools and BCFtools. GigaScience 10, giab008 (2021).

65. McKenna, A. et al. The Genome Analysis Toolkit: A MapReduce framework for analyzing next-generation DNA sequencing data. Genome Res. 20, 1297–1303 (2010).

66. Chen, X. et al. Manta: rapid detection of structural variants and indels for germline and cancer sequencing applications. Bioinformatics 32, 1220–1222 (2016).

67. Saunders, C. T. et al. Strelka: accurate somatic small-variant calling from sequenced tumor–normal sample pairs. Bioinformatics 28, 1811–1817 (2012).

68. Park, J. et al. Accurate somatic small variant discovery for multiple sequencing technologies with DeepSomatic. Nat Biotechnol 44, 1569–1578 (2026).

69. Pablo Riesgo-Ferreiro, Özlem Muslu & Luis Kress. TRON-Bioinformatics/vafator: vafator v3.0.0. Zenodo 10.5281/ZENODO.19497466 (2026).

70. McLaren, W. et al. The Ensembl Variant Effect Predictor. Genome Biol 17, 122 (2016).

71. Jaganathan, K. et al. Predicting Splicing from Primary Sequence with Deep Learning. Cell 176, 535–548.e24 (2019).

72. Liu, X., Li, C., Mou, C., Dong, Y. & Tu, Y. dbNSFP v4: a comprehensive database of transcript-specific functional predictions and annotations for human nonsynonymous and splice-site SNVs. Genome Med 12, 103 (2020).

73. Köster, J., Dijkstra, L. J., Marschall, T. & Schönhuth, A. Varlociraptor: enhancing sensitivity and controlling false discovery rate in somatic indel discovery. Genome Biol 21, 98 (2020).

74. Hao, Y. et al. Dictionary learning for integrative, multimodal and scalable single-cell analysis. Nat Biotechnol 42, 293–304 (2024).

75. Lareau, C. A. et al. Massively parallel single-cell mitochondrial DNA genotyping and chromatin profiling. Nat Biotechnol 39, 451–461 (2021).

76. Kuipers, J., Jahn, K., Raphael, B. J. & Beerenwinkel, N. Single-cell sequencing data reveal widespread recurrence and loss of mutational hits in the life histories of tumors. Genome Res 27, 1885–1894 (2017).

77. Yu, G., Smith, D. K., Zhu, H., Guan, Y. & Lam, T. T. GGTREE : an R package for visualization and annotation of phylogenetic trees with their covariates and other associated data. Methods Ecol Evol 8, 28–36 (2017).

78. Ludwig, L. S. et al. Lineage Tracing in Humans Enabled by Mitochondrial Mutations and Single-Cell Genomics. Cell 176, 1325–1339.e22 (2019).

79. Stuart, T., Srivastava, A., Madad, S., Lareau, C. A. & Satija, R. Single-cell chromatin state analysis with Signac. Nat Methods 18, 1333–1341 (2021).

80. Huang, Y., McCarthy, D. J. & Stegle, O. Vireo: Bayesian demultiplexing of pooled single-cell RNA-seq data without genotype reference. Genome Biol 20, 273 (2019).

81. Thibodeau, A. et al. AMULET: a novel read count-based method for effective multiplet detection from single nucleus ATAC-seq data. Genome Biol 22, 252 (2021).

82. Zhang, Y. et al. Model-based Analysis of ChIP-Seq (MACS). Genome Biol 9, R137 (2008).

83. Li, H. & Durbin, R. Fast and accurate short read alignment with Burrows–Wheeler transform. Bioinformatics 25, 1754–1760 (2009).

84. Li, H. et al. The Sequence Alignment/Map format and SAMtools. Bioinformatics 25, 2078–2079 (2009).

85. Wang, K., Li, M. & Hakonarson, H. ANNOVAR: functional annotation of genetic variants from high-throughput sequencing data. Nucleic Acids Research 38, e164–e164 (2010).

86. Quinlan, A. R. & Hall, I. M. BEDTools: a flexible suite of utilities for comparing genomic features. Bioinformatics 26, 841–842 (2010).

87. Bergstrom, E. N. et al. SigProfilerMatrixGenerator: a tool for visualizing and exploring patterns of small mutational events. BMC Genomics 20, 685 (2019).

88. Islam, S. M. A. et al. Uncovering novel mutational signatures by de novo extraction with SigProfilerExtractor. Cell Genomics 2, 100179 (2022).

89. Durrett, R. Probability Models for DNA Sequence Evolution. (Springer, New York ; London, 2008).

90. Fodil-Cornu, N., et al. *Ly49h* -Deficient C57BL/6 Mice: A New Mouse Cytomegalovirus-Susceptible Model Remains Resistant to Unrelated Pathogens Controlled by the NK Gene Complex. The Journal of Immunology 181, 6394–6405 (2008).

91. Lee, J.-E. et al. H3K4 mono- and di-methyltransferase MLL4 is required for enhancer activation during cell differentiation. Elife 2, e01503 (2013).

